# Molecular basis for gating of cardiac ryanodine receptor: underlying mechanisms for gain- and loss-of function mutations

**DOI:** 10.1101/2020.11.30.401026

**Authors:** Takuya Kobayashi, Akihisa Tsutsumi, Nagomi Kurebayashi, Kei Saito, Masami Kodama, Takashi Sakurai, Masahide Kikkawa, Takashi Murayama, Haruo Ogawa

## Abstract

Cardiac ryanodine receptor (RyR2) is a large Ca^2+^ release channel in the sarcoplasmic reticulum and indispensable for excitation-contraction coupling in the heart. RyR2 is activated by Ca^2+^ and RyR2 mutations are implicated in severe arrhythmogenic diseases. Yet, the structural basis underlying channel opening and how mutations affect the channel remain unknown. Here, we addressed gating mechanism of RyR2 by combining high-resolution structures determined by cryo-electron microscopy with quantitative functional analysis of channels carrying various mutations in specific residues. We demonstrated two fundamental mechanisms for channel gating: interactions close to the channel pore stabilize the channel to prevent hyperactivity and a series of interactions in the surrounding regions is necessary for channel opening upon Ca^2+^ binding. Mutations at the residues involved in the former and the latter mechanisms cause gain-of-function and loss-of-function, respectively. Our results reveal gating mechanisms of the RyR2 channel and alterations by pathogenic mutations at the atomic level.

## Introduction

Cardiac ryanodine receptor (RyR2) is a Ca^2+^ release channel in the sarcoplasmic reticulum and plays a central role in cardiac muscle contraction (Bers, 2004; Santulli et al., 2018). In cardiac excitation-contraction coupling, Ca^2+^ influx occurred through L-type voltage-dependent Ca^2+^ channels in the transverse (T) tubule membrane activates RyR2 to release large amount of Ca^2+^, a process known as Ca^2+^-induced Ca^2+^ release (CICR) (Endo, 2009; Fabiato and Fabiato, 1978). In human RyR2, nearly 300 pathogenic mutations have been reported as arrhythmogenic heart diseases, including catecholaminergic polymorphic ventricular tachycardia (CPVT) (Kawamura et al., 2013; Pérez-Riera et al., 2018; Priori et al., 2002; Tester et al., 2004), and idiopathic ventricular fibrillation (IVF) (Fujii et al., 2017; Hirose et al., 2021; Jiang et al., 2007; Paech et al., 2014). Notably, these mutations divergently alter channel activity: mutations related to CPVT cause gain-of-function, whereas those for IVF lead to either gain- or loss-of-function (Benkusky et al., 2004; Priori and Chen, 2011).

RyR is a large tetrameric ion channel (∼2.2 MDa), with each monomer composed of ∼5,000 amino acid residues and comprising a large N-terminal mushroom-like structure of the cytoplasmic domain and C-terminal transmembrane (TM) region (Clarke and Hendrickson, 2016; Gong et al., 2021; Zalk and Marks, 2017). The opening of RyR channels is initiated by binding of Ca^2+^ to the Ca^2+^-binding site, which was proposed at the interface between Central and CTD domains by structural analysis using cryo-electron microscopy (EM) (des Georges et al., 2016) and, later, validated by functional analysis using site-directed mutagenesis (Murayama et al., 2016; Xu et al., 2018). The mechanism of channel opening after Ca^2+^ binding has been extensively studied by structural analysis of the channel at the open state (Bai et al., 2016; Chi et al., 2019; des Georges et al., 2016; Gong et al., 2019; Peng et al., 2016; Wei et al., 2016). These results demonstrated that Ca^2+^-binding causes a large conformational change throughout the molecule via a large number of intra/inter-domain interactions, thereby the channel opens.

The activity of the RyR channels is modulated by various small molecules and associated proteins (Meissner, 2017). The RyR structures in complex with regulatory molecules, such as ATP, caffeine, FKBP12 or calmodulin, have also been determined, and the mechanisms by which these regulatory molecules modulate RyR have been extensively discussed (Chi et al., 2019; des Georges et al., 2016; Gong et al., 2019). In addition, the structures of three different gain-of-function mutants with mutations in the N-terminal domains have recently been reported (Iyer et al., 2020; Woll et al., 2021). In all structures, slight slippages in the inter-domain interactions around the mutated site occur, and both groups concluded that these slippages may lead to the increase in the open probability in the channel through inter-domain interactions.

Although many structures of RyR have been so far determined, the molecular mechanism of channel gating of RyR still remains largely obscured, due to lack of our knowledge about the key interactions for the conformational change, and how and in what order these multiple interaction-networks interlock for the channel opening. In addition, all the open structures obtained to date contain additional ligands (ATP, caffeine, and/or PCB95), which help opening of the channel but might make it difficult to discriminate conformational changes by Ca^2+^. To precisely trace the Ca^2+^-induced conformational changes, it is necessary to obtain open structure with Ca^2+^ alone.

In this study, we newly determined high-resolution closed and open structures of recombinant mouse RyR2, in which open structure was successfully obtained by Ca^2+^ alone. Then, we performed extensive functional analysis of the RyR2 channels carrying mutations in amino acid residues in the proposed interactions. Finally, structures of a loss-of-function mutant were determined to prove our hypothesis of gating mechanism. We demonstrated two fundamental mechanisms for channel gating of RyR2. In the resting state without Ca^2+^, the channel is stabilized in the closed state by multiple interactions close to the channel pore. Upon Ca^2+^ binding, a series of interactions in the surrounding regions moves the S4-S5 linker outward to open the gate. Disruption of the former and the latter interactions causes gain-of-function and loss-of-function, respectively. Our results reveal mechanisms underlying channel opening upon Ca^2+^ binding and explain how pathogenic mutations alter channel activity.

## RESULTS

### Overall conformational changes in RyR2 associated with Ca^2+^ binding

Recombinant mouse RyR2, expressed and purified from HEK293 cells using FKBP12.6 affinity chromatography (Cabra et al., 2016) formed homogeneous tetrameric channels (Figures. S1A and S1B). To precisely trace the Ca^2+^-induced RyR2 structural changes, we adopted high salt conditions, which markedly enhance channel activity with Ca^2+^ alone (Figures S1C–S1E) (Meissner, 1994; Ogawa, 1994). We performed cryo-EM single-particle analysis in the presence of 1 mM EGTA and 100 μM Ca^2+^ for closed and open states, respectively. Three-dimensional (3D) classification including focused classification analysis using the TM region revealed two and three classes in the presence of EGTA and Ca^2+^, respectively (Figures S1F and S1G; Table S1). No major differences were observed among the classes both in the closed and open states. Therefore, subsequent analysis was performed using the class with the highest resolution (before the final classification in the closed state and class 1 in the open state; Table S1). The overall resolution of the final model was 3.3 Å and 3.45 Å for the closed and open states, respectively, and the local resolution for the TM region in the closed state was better than 2.9 Å (Figures S1F and S1G), which allowed us to build precise atomic models and identify specific residues important for channel gating (Figures S1H and S1I). In fact, it was confirmed that the pore was open only by addition of 100 μM of Ca^2+^ (Figure S2A), and moreover, the bound FKBP12.6 is clearly visible in both the closed and open states (Figures S1H and S1I). Conformational changes between closed and open states were large and spanned the entire molecule (Figures 1A, 1B, S2B-S2D; Video S1). Changes associated with Ca^2+^ binding were analyzed in three layers with different heights parallel to the membrane, i.e., the C-terminal domain (CTD), U-motif, and S4-S5 layers (Figures 1C–1E; Video S1). Upon Ca^2+^ binding, CTD rotated clockwise (viewed from cytoplasmic side) (Figure 1C), U-motif rotated clockwise toward the S2-S3 domain (Figure 1D), and S6 leaned outward from the center to open the gate accompanied by rearrangements of S1-S4 helices and outward movement of the S4-S5 linker (Figure 1E). The C-terminal side of the Central domain (C-Central), U-motif, CTD, and cytoplasmic S6 (S6_cyto_) were tightly attached to each other in both closed and open states (Figures 1F, S2E, and S2F; Video S2) and rotated together 9.8° clockwise with respect to the axis approximately parallel to S6 (Figure 1G). In contrast, the N-terminal side of the Central domain (N-Central) did not follow the rotation due to splitting the Central domain into two parts with G3987 as the pivot (Figure 1F). Since the rotation axis was close to the Ca^2+^-binding site, slight movements caused by Ca^2+^ binding resulted in very large movements around S6 (Figure 1G).

**Fig. 1.**
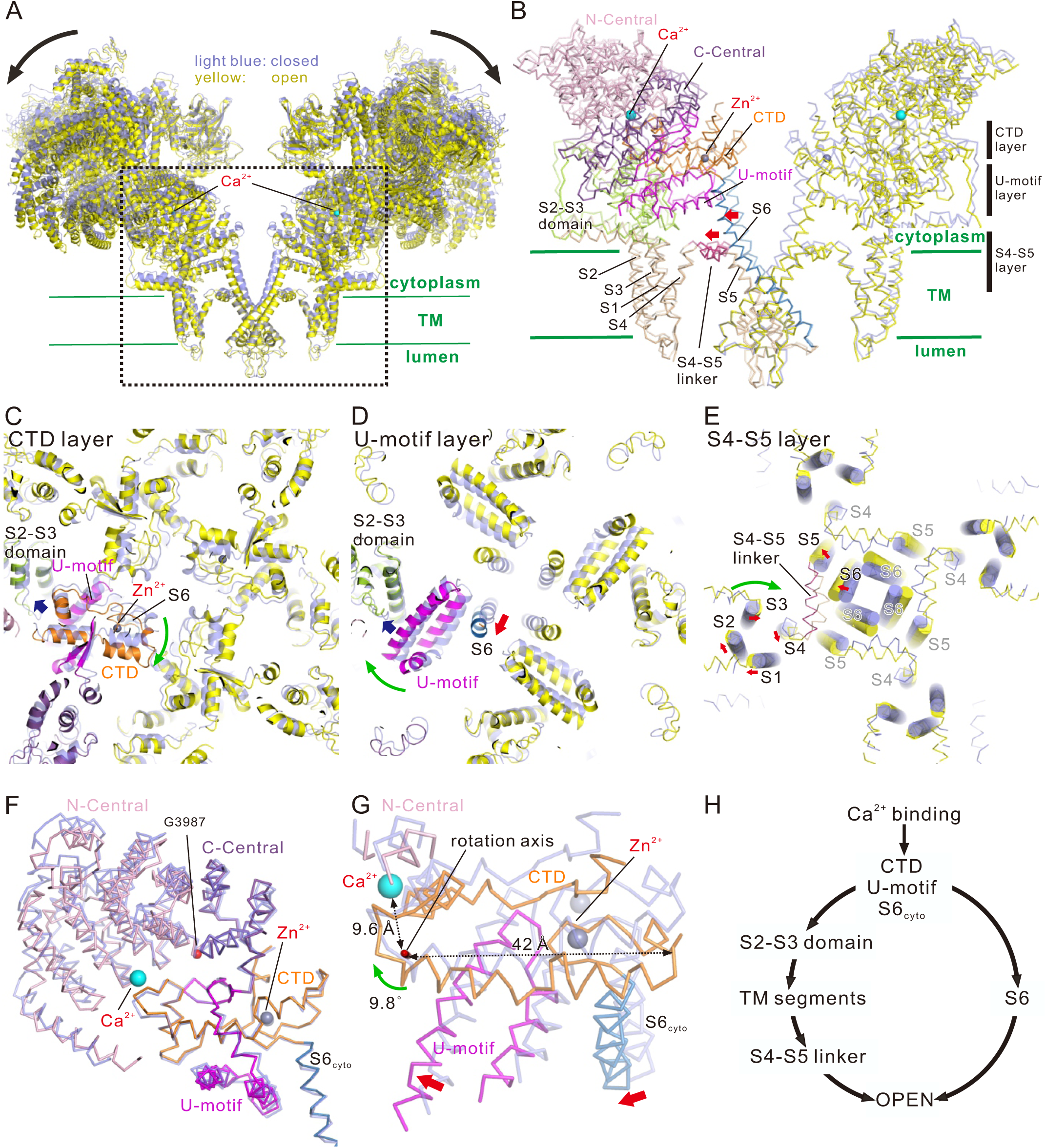
Conformational changes upon Ca^2+^ binding. (A) Overlay of RyR2 in the closed (light blue) and open (yellow) states viewed from the direction parallel to the lipid bilayer is shown as a ribbon model. Two facing protomers in the RyR2 tetramer are shown. (B) Magnified view of the dotted box in (A). In the left protomer, each domain is colored (N-Central, light pink; C-Central, purple; U-motif, magenta; S1-S5, wheat; S2-S3 domain, light green; S4-S5 linker, warm pink; S6, blue; CTD, orange). S4-S5 linker and S6 moved outside upon Ca^2+^ binding as indicated by the red arrows. Three regions parallel to the membrane are defined as CTD, U-motif, and S4-S5 layers. Ca^2+^, shown as cyan ball; Zn^2+^, shown as gray ball. (C–E) Cross-section views of CTD, U-motif, and S4-S5 layers. Closed state is colored in light blue and open state is colored according to (B) or yellow. In (E), Cα representation overlaid with cylindrical TM helices are used. Ca^2+^ binding causes clockwise rotation of CTD (green arrow in C), U-motif (green arrow in D), and S1-S4 TM helices and outward movement of S4-S5 linker and S6 (red arrows in E). (F) C-Central/U-motif/S6_cyto_/CTD complex. Closed (light blue) and open (colored according to B) states are overlaid at the CTD. Central domain is split into two parts at G3987 which works as the pivot of the rotation upon Ca^2+^ binding. (G) Rotation of the C-Central/U-motif/S6_cyto_/CTD complex upon Ca^2+^ binding viewed from the rotation axis. (H) Scheme of channel opening upon Ca^2+^ binding. Two independent pathways via S6 and S4-S5 linker are hypothesized.

Given these series of movements, we hypothesized that the rotation of the C-Central/U-motif/S6_cyto_/CTD complex upon Ca^2+^ binding leads to two independent downstream pathways for channel pore opening; (1) the S6_cyto_ movement leads to the outward leaning of S6; (2) the U-motif movement leads to sequential movements of the S2-S3 domain and TM segments, causing outward movement of the S4-S5 linker and creating a space where S6 can lean into (Figure 1H). To prove this hypothesis, we performed detailed structural analysis and functional studies.

### U-motif/S2-S3 domain interaction is a key for signal transduction in response to Ca^2+^ binding

In our hypothesis, the U-motif/S2-S3 domain interaction is important in transducing the rotation of the U-motif to TM segment movement (Figure 1D; Videos S1 and S2). We found three key interactions between the U-motif and S2-S3 domain formed by hydrogen bonds or salt bridges (Figures 2A–2C and S3A; Video S3). E4198-K4593-S4167 was evident in the open state, whereas Y4498-K4593 was only formed in the closed state, indicating that K4593 switches the interacting partner between the two states. Additionally, E4193-R4607 was stable in both states.

**Fig. 2.**
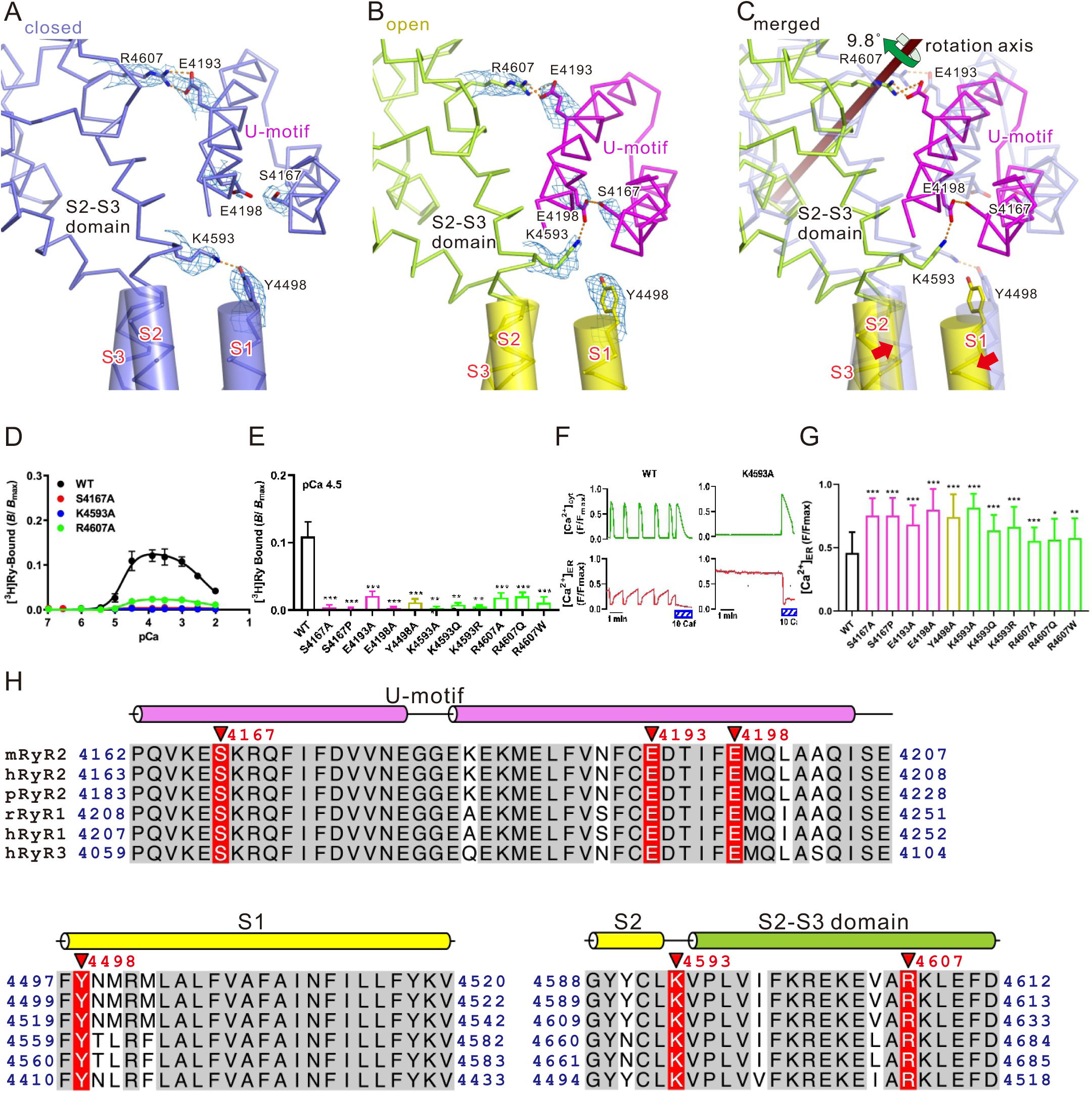
Key interactions between U-motif and S2-S3 domain upon channel opening. (A–C) Interface of U-motif and S2-S3 domain in the closed state (A), open state (B), and an overlay of both states (C) viewed parallel to the membrane is shown as a Cα model. Amino acid residues involved in key interactions are shown as stick models. The color of carbon atoms is the same as that of Cα; oxygen, red; nitrogen, blue. The TM region forming α-helices is overlaid with the cylinder model. Hydrogen bonds or salt bridges are shown as orange dotted lines. Density maps around side chains shown in (A) and (B) are superimposed and contoured at 0.025. (D–G) Functional analysis of mutants involved in U-motif/S2-S3 domain interaction. (D) Ca^2+^-dependent [^3^H]ryanodine binding of WT and representative mutants. (E) Summary of [^3^H]ryanodine binding of WT and mutants at pCa 4.5. (F) Representative traces of cytoplasmic ([Ca^2+^]_cyt_) and ER ([Ca^2+^]_ER_) Ca^2+^ signals of HEK293 cells expressing WT or K4593A. Spontaneous Ca^2+^ oscillations occurred with a concomitant decrease in [Ca^2+^]_ER_ in WT, while the K4593 mutant showed no Ca^2+^ oscillations with an increased [Ca^2+^]_ER_, indicating loss-of-function of the channel. (G) Summary of upper level of [Ca^2+^]_ER_ signals in WT and mutants. All mutants showed loss-of-function of the channel. Data are presented as the mean ± SD. *p < 0.05; **p < 0.01; ***p < 0.001. (H) Multiple sequence alignment of three RyR isoforms around the U-motif, S1, and S2 to the initial S2-S3 domain. The gray-shaded residues are the identical sequences among isoforms, and the residues shown in red-shade are the residues for which the mutants were prepared and the functional assays were performed. Secondary structures are shown above the alignment. m, mouse; h, human; p, pig; r, rabbit.

To clarify the roles of hydrogen bonds/salt bridges in channel gating, functional assays were conducted with recombinant RyR2 carrying mutations at the specific residues (Table 1). Ca^2+^-dependent [^3^H]ryanodine binding was conducted as it reflects channel activity (Fujii et al., 2017; Murayama et al., 2018b). Wild-type (WT) RyR2 exhibited biphasic Ca^2+^-dependent [^3^H]ryanodine binding, whereas alanine substitution at K4593 markedly reduced the binding (Figure 2D). Two pathogenic mutations, K4593Q and K4593R (Hirose et al., 2021), also led to loss of binding (Figures 2E and S3B). Similarly, binding was severely reduced after alanine substitutions (Y4498A, S4167A, and E4198A) and pathogenic mutation (S4167P) (Hirose et al., 2021) in interacting partners (Figures 2D, 2E and S3B). The behaviors of mutant RyR2 channels were also evaluated in live cells by monitoring cytoplasmic and endoplasmic reticulum (ER) luminal Ca^2+^ homeostasis (Fujii et al., 2017; Uehara et al., 2017). In WT RyR2-expressing HEK293 cells, spontaneous Ca^2+^ oscillations occurred with a concomitant decrease in [Ca^2+^]_ER_, indicating Ca^2+^ release from the ER via RyR2 channels (Figure 2F, left). In contrast, RyR2 channels carrying the K4593A mutation showed no such Ca^2+^ oscillations with increased [Ca^2+^]_ER_, indicating a loss-of-function of the channel (Figure 2F, right). Similar results were obtained with other substitutions in S4167, E4198, and K4593 (Figure 2G). Altogether, these results suggest that both E4198-K4593-S4167 and Y4498-K4593 interactions are important for channel opening. We also evaluated the E4193-R4607 interaction and found that alanine substitutions (E4193A and R4607A) and pathogenic mutations (R4607Q and R4607W) led to a loss-of-function of the channel (Figures 2D–2G, S3B). These residues and surrounding sequences are well conserved in all three mammalian RyR subtypes (Figure 2H). These findings indicate that three interactions above S2 in the U-motif/S2-S3 domain interface play a critical role in transducing the Ca^2+^-binding signal to S2, and that loss of these interactions results in a loss of channel function.

**Table 1.** RyR2 mutants used in this study.

| **Mutation** | **Domain** | **Disease** | **Ref** | **Ryanodine binding** | **ER Ca<sup>2+</sup> level** | **Functional change** |
| --- | --- | --- | --- | --- | --- | --- |
| S4167A | U-motif |  |  | ↓ | ↑ | LOF |
| S4167P | U-motif | CPVT, LQTS (RyR2) | (Ozawa et al., 2018) | ↓ | ↑ | LOF |
| F4171A | U-motif |  |  | ↑ | ↓ | GOF |
| I4172A | U-motif |  |  | ↑ | ↓ | GOF |
| F4173A | U-motif |  |  | ↑ | ↓ | GOF |
| V4175A | U-motif |  |  | ↑ | ↓ | GOF |
| V4176A | U-motif |  |  | ↑ | ↓ | GOF |
| N4177A | U-motif |  |  | ↑ | ↓ | GOF |
| N4177S | U-motif | CPVT (RyR2) | (Medeiros-Domingo et al., 2009) | ↑ | ↓ | GOF |
| N4177Y | U-motif | CPVT (RyR2) | (Hayashi et al., 2009) | ↑ | ↓ | GOF |
| F4191A | U-motif |  |  | ↓ |  | GOF |
| E4193A | U-motif |  |  | ↓ | ↑ | LOF |
| E4198A | U-motif |  |  | ↓ | ↑ | LOF |
| F4497A | S1 |  |  | ↓ | ↑ | LOF |
| F4497C | S1 | CPVT (RyR2) | (Choi et al., 2004) | ↓ | ↑ | LOF |
| Y4498A | S1 |  |  | ↓ | ↑ | LOF |
| R4501A | S1 |  |  | ↑ | ↓ | GOF |
| L4505A | S1 |  |  | ↑ | ↓ | GOF |
| L4505P | S1 | CCD (RyR1) | (Wu et al., 2006) | ↑ | ↓ | GOF |
| Y4589A | S2 |  |  | ↓ | ↑ | LOF |
| L4592A | S2 |  |  | ↓ | ↑ | LOF |
| K4593A | S2-S3 |  |  | ↓ | ↑ | LOF |
| K4593Q | S2-S3 | LQTS (RyR2) | (Hirose et al., 2021; Shigemizu et al., 2015) | ↓ | ↑ | LOF |
| K4593R | S2-S3 | IVF (RyR2) | (Paech et al., 2014) | ↓ | ↑ | LOF |
| R4607A | S2-S3 |  |  | ↓ | ↑ | LOF |
| R4607Q | S2-S3 | Sudden death (RyR2) | (Wong et al., 2014) | ↓ | ↑ | LOF |
| R4607W | S2-S3 | Sudden death (RyR2) | (Wang et al., 2014) | ↓ | ↑ | LOF |
| D4715A | S3 |  |  | ↓ | ↑ | LOF |
| Y4720A | S3 |  |  | ↑ | ↓ | GOF |
| Y4720C | S3 | CPVT (RyR2) | (Roston et al., 2018) | ↑ | ↓ | GOF |
| D4744A | S4 |  |  | ↑ | ↓ | GOF |
| D4744H | S4 | Myopathy (RyR1) | (Bharucha-Goebel et al., 2013) | ↑ | ↓ | GOF |
| F4749A | S4 |  |  | ↑ | ↓ | GOF |
| F4749V | S4 |  |  | ↓ |  | LOF |
| Q4875A | S6cyto |  |  | ↑ | ↓ | GOF |
| V4879A | S6cyto | CPVT (RyR2) | (Bagattin et al., 2004) | ↑ | ↓ | GOF |
| F4888A | CTD |  |  | ↑ | ↓ | **GOF** |
| F4888Y | CTD | MH/CCD (RyR1) | (Ibarra M et al., 2006) | ↑ | ↓ | **GOF** |
| L4914A | CTD |  |  | ↑ | ↓ | **GOF** |

CTD, C-terminal domain; CPVT, catecholaminergic polymorphic ventricular tachycardia; CCD, central core disease; LQTS, long QT syndrome; IVF, idiopathic ventricular fibrillation; MH, malignant hyperthermia; LOF, loss-of-function; GOF, gain-of-function.

### Movements of the S1-S4 bundle lead to outward movement of the S4-S5 linker

Movement of the S2-S3 domain appears to cause S2 movement (see Figure 2C and video S3). This leads to the coordinated 7.6° clockwise rotation of S1, S3, and S4 (Figures 1E, 3A–3C, S4A-S4C; Videos S1 and S4). S1, S2, S3, and S4 are arranged in a circle, placed at equal intervals in a clockwise direction (Figures 3A–3C, S4A-S4C). We found interactions between S1 and S2 (S1/S2), S2/S3, S3/S4, and S1/S4, all of which were maintained in both closed and open states (Figures 3A and 3B, S4A-S4C; Video S4). The hydrophobic interactions between F4497 and L4592 at S1/S2; hydrogen bond between Y4589 and D4715 at S2/S3; hydrogen bond between Y4720 and D4744 at S3/S4; and the salt bridge between R4501 and D4744 at S1/S4 appear to bundle the four TM segments into one (i.e., S1-S4 bundle).

**Fig. 3.**
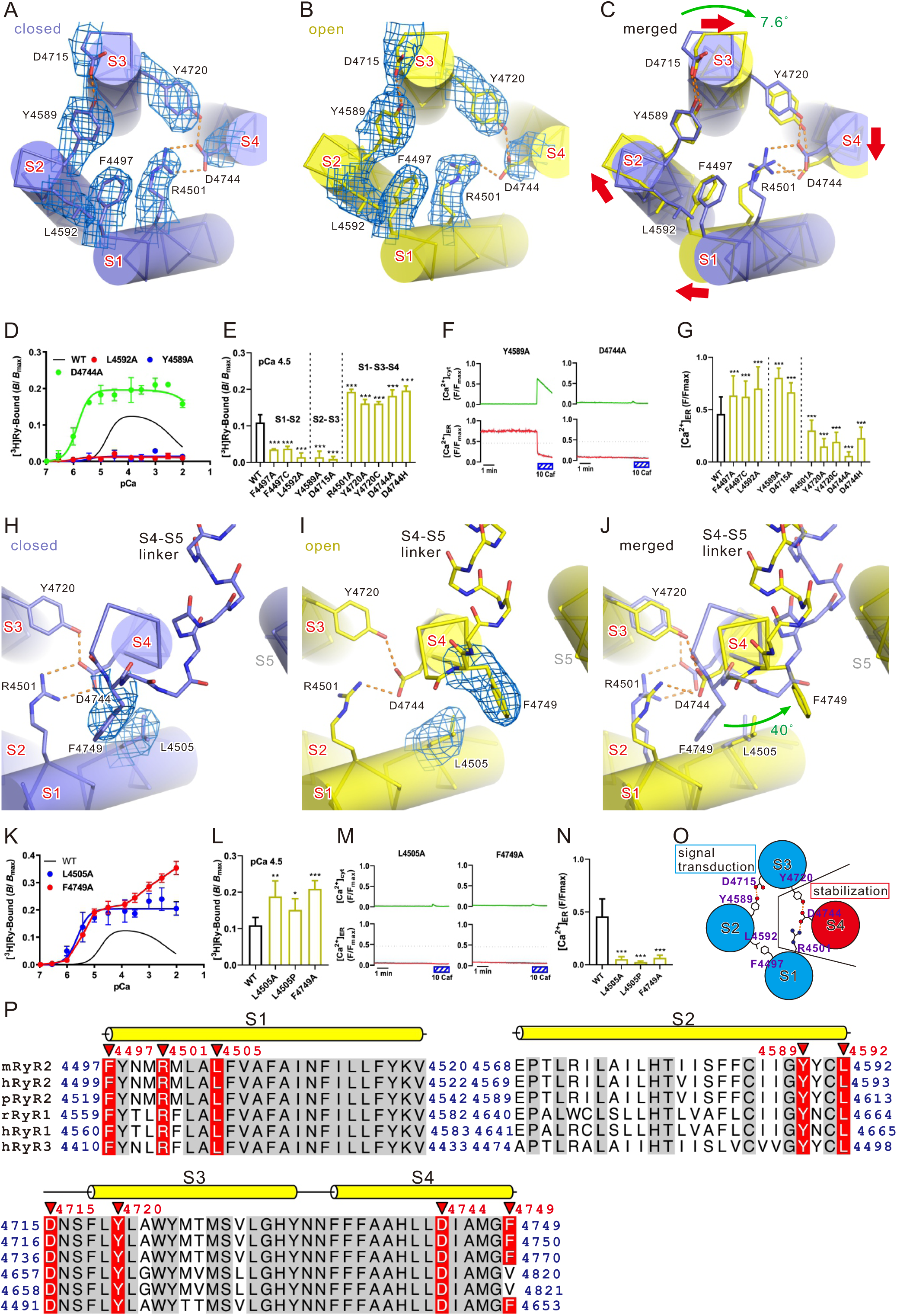
Key interactions in the transmembrane region upon channel opening. (A–C) The S1-S4 bundle. Closed state (A), open state (B), and overlay of the structures in both states (C) are shown as a Cα model and overlaid with cylinder models. Hydrogen bonds/salt bridges are shown as orange botted lines. Density maps around side chains shown in (A) and (B) are superimposed and contoured at 0.03. (D–G) Functional analysis of mutants involved in the S1/S2, S2/S3, S1/S4, or S3/S4 interaction. (D) Ca^2+^-dependent [^3^H]ryanodine binding of WT and representative mutants. (E) Summary of [^3^H]ryanodine binding of WT and mutants at pCa 4.5. (F) Representative traces of [Ca^2+^]_cyt_ and [Ca^2+^]_ER_ signals of HEK293 cells expressing Y4589A or D4744A. (G) Summary of upper level of [Ca^2+^]_ER_ signals in WT and mutants. (H–J) TM region around S4. Closed state (H), open state (I), and overlay of the structures in both states (J). Structures in both states were fitted to the bottom part of S4 helices and viewed along the axis of the S4 helix and from the cytoplasm; Cα model overlaid with cylinder models. Main chain representation of the S4-S5 linker. Density maps around side chains shown in (H) and (I) are superimposed and contoured at 0.03. (K) Ca^2+^-dependent [^3^H]ryanodine binding of WT and representative mutants. (L) Summary of [^3^H]ryanodine binding of WT and mutants at pCa 4.5. (M) Representative traces of [Ca^2+^]_cyt_ and [Ca^2+^]_ER_ signals of HEK293 cells expressing L4505A or F4749A. (N) Summary of upper level of [Ca^2+^]_ER_ signals in WT and mutants. (O) Scheme of the structure in the S1-S4 TM helices. While the interactions between S1 and S2 or S2 and S3 are involved in signal transduction, the interactions between S1 and S4 or S3 and S4 are involved in the stabilization of the channel in the closed state. (P) Multiple sequence alignment of three RyR isoforms around S1, S2, and S3 to S4. The gray-shaded residues are the identical sequences among isoforms, and the residues for which the mutants were prepared and the functional assays were performed. Secondary structures are shown above the alignment. m, mouse; h, human; p, pig; r, rabbit.

Alanine substitution of residues involved in S1/S2 (F4497 and L4592) and S2/S3 (Y4589 and D4715) interactions exhibited significantly reduced [^3^H]ryanodine binding (Figures 3D, 3E, and S4G), loss of Ca^2+^ oscillations, and increased ER Ca^2+^ (Figures 3F and 3G)—all indicative of loss-of-function. The pathogenic mutant F4497C also resulted in loss-of-function (Figures 3E, 3G and S4G). Thus, S1/S2 and S2/S3 interactions are necessary for channel opening. In contrast, alanine substitution or pathogenic mutations of residues involved in S1/S4 (R4501 and D4744) and S3/S4 (Y4720 and D4744) interactions caused gain-of-function of the channel with increased [^3^H]ryanodine binding and reduction in ER Ca^2+^ (Figures 3D-3G, and S4G). Especially, D4744A, which is involved in both interactions, exhibited greatly increased [^3^H]ryanodine binding with an enhanced Ca^2+^ sensitivity for activation and loss of Ca^2+^ inactivation (Figures 3D, S4I, and S4J).

Rotation of the S1-S4 bundle causes movement of the S4-S5 linker. We found that the upper part of S4 rotated 40° counterclockwise to form an α-helix upon channel opening, which dramatically changed the position of F4749 (Figures 3H–3J and S4A-S4F; Video S4). The S4-S5 linker, which is unfolded and significantly bent in the direction of S6 in the closed state, rewinds to an α-helix and moves outward (Figures S4D-S4F; Video S4). A hydrophobic interaction between F4749 in the upper part of S4 and L4505 in S1 was identified in the closed state (Figures 3H and S4A).

Alanine substitutions of residues involved in hydrophobic interactions (L4505A and F4749A) and the pathogenic mutant L4505P caused gain-of-function with increased Ca^2+^ sensitivity and loss of Ca^2+^ inactivation (Figures 3K–3N, S4H-S4J). Notably, no additive effects were observed upon binding of [^3^H]ryanodine to double mutant L4505A_F4749A (Figure S4H), supporting an interaction between the two residues. Whereas L4505 is conserved in all the mammalian RyR subtypes, F4749 is mutated to valine in the RyR1 subtype (Figure 3O). However, unlike alanine mutant, F4749V exhibited biphasic Ca^2+^ dependent [^3^H]ryanodine binding with slightly reduced peak value (Figures S4H). Brief summary of the roles of the key residues located on the S1-S4 bundle is shown as a scheme (Figure 3O). These residues are well conserved in all the mammalian RyR subtypes (Fig. 3P).

Taking these findings into consideration, we hypothesized a channel opening mechanism in which a series of movements of S4-S5 linker regulate channel opening. In the absence of Ca^2+^, the channel is stabilized in the closed state as the S4-S5 linker is locked by the “stopper,” in form of L4505-F4749 interaction, which prevents α-helix formation of the upper part of S4. The L4505-F4749 interaction is supported by S1/S3 and S3/S4 interactions that keep the two residues appropriately placed. Upon Ca^2+^ binding, clockwise rotation of the S1-S4 bundle induced by U-motif/S2-S3 domain interaction alters the relative positions of L4505 and F4749 to release the stopper. This allows α-helix formation of the upper part of S4 and subsequent outward movement of the S4-S5 linker to open the channel. Loss of necessary interactions, such as that of the U-motif/S2-S3 domain (Figures 2A–2C), S1/S2, and S2/S3 (Figures 3A–3C), cannot release the stopper, leading to loss-of-function (Figures 2D–2G and 3D–3G). Conversely, loss of interactions that support S4, such as S1/S4 and S3/S4 (Figures 3A–3C and 3H–3J) destabilize the channel in the closed state and impart a gain-of-function of the channel (Figures 3D–3G and 3K–3N).

### U-motif plays key role in stabilizing the channel in the closed state

U-motif constitutes the U-motif/S6_cyto_/CTD complex, which was stable in both the closed and open states (Figure 1F). We found a close contact between U-motif and S6_cyto_ as observed in the closed state structure of RyR1 (Yan et al., 2015) (Figures 4A, 4B, and S5A; Video S5). In the closed state, F4173, V4176, and N4177 in the U-motif and Q4875 and V4879 in S6_cyto_ faced each other forming van der Waals interactions, and were surrounded by Q4876 and Q4878 from S6 (Figure 4B; Video S5). In the open state, S6_cyto_ self-rotated (∼30º clockwise at Q4875) and U-motif/S6_cyto_ loosened and appeared unstable (Figure 4C and S5B; Video S5). Buried surface area analysis calculated by CNS (Brünger et al., 1998) demonstrated that the U-motif/S6_cyto_ interface in the open state (362 Å^2^) was smaller than in the closed state (514 Å^2^). The U-motif also interacted with CTD via F4888, where it penetrated partly into the hydrophobic pocket formed by U-motif F4171, I4172, and V4175 and CTD L4914 (Figure 4D and S5C; Video S5). The U-motif/CTD interaction was stable in both states.

**Fig. 4.**
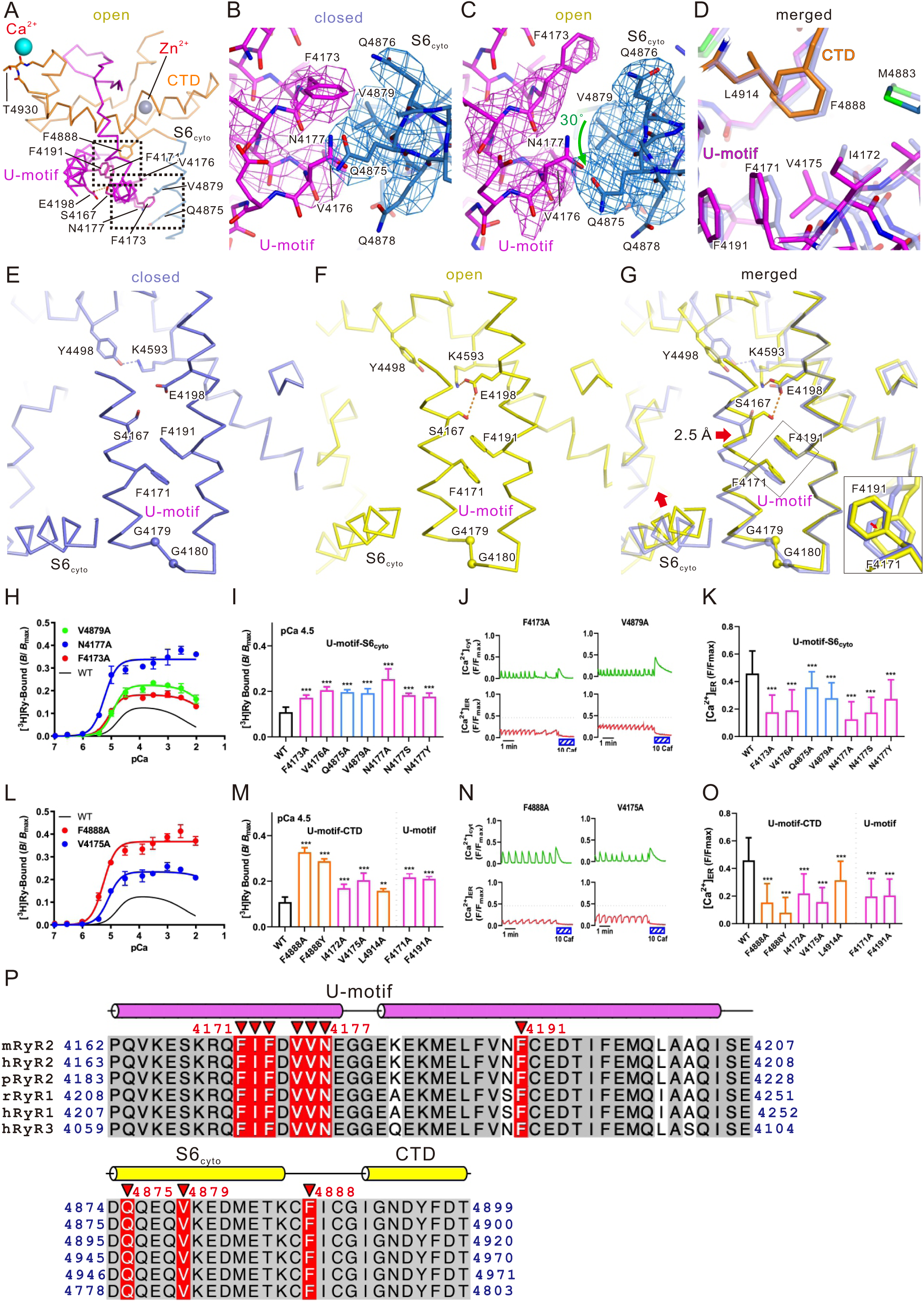
Key interactions between the U-motif and S6_cyto_/CTD. (A) Structure around the U-motif in the open state (U-motif, magenta; S6_cyto_, blue; CTD, orange). Ca^2+^ and Zn^2+^ are shown as cyan and gray spheres, respectively. (B, C) Details of the U-motif/S6_cyto_ interaction. Structures in the closed (B) and open (C) states fitted to the N-terminal region of the U-motif are shown as a full atomic model. Density maps around the interaction are superimposed and contoured at 0.025. (D) Details of the U-motif/CTD interaction around F4888. Overlay of the structures in the closed (light blue) and open (colored) states. (E–G) Compactization in U-motif. Closed state (E), open state (F), and overlay of the structures in both states (G) are shown as a Cα model. Inset of (G) shows the magnified view of the F4171/F4191 stacking looking from aromatic ring of F4191. F4171 is stacked in parallel with F4191 in both the closed state and the open state, but it moves a little away from F4191 in the open state. The structures are fitted in the C-terminal side of U-motif (4183-4205). Closed state and open state is colored with light blue and yellow, respectively. The N-terminus side of the U-motif of the WT in the open state is ∼2.5 Å closer to the C-terminus side of the U-motif as indicated by the red arrow, and as a result, S6_cyto_ movement (red arrow) becomes possible. (H–K) Functional analysis of mutants involved in U-motif/S6_cyto_ (E–H) and U-motif/CTD (L–O) interactions. (H, L) Ca^2+^-dependent [^3^H]ryanodine binding of WT and representative mutants. (I, M) Summary of [^3^H]ryanodine binding of WT and mutants at pCa 4.5. (J, N) Representative traces of [Ca^2+^]_cyt_ and [Ca^2+^]_ER_ signals of HEK293 cells expressing F4173A or V4879A (J) and F4888A or V4175A (N). (K, O) Summary of upper level of [Ca^2+^]_ER_ signals in WT and mutants. (P) Multiple sequence alignment of three RyR isoforms around the U-motif and S6_cyto_ to the CTD. The gray-shaded residues are the identical sequences among isoforms, and the residues in which mutants were prepared and functional assayed are red-shaded. Secondary structures are shown above the alignment. m, mouse; h, human; p, pig; r, rabbit.

We also noticed compaction of U-motif upon channel opening, which was induced by a 2.5 Å parallel shift of its N-terminal helix (4161-4178) toward the C-terminal side (4183-4205) (Figures 4E-4G). This parallel shift seems to be the key trigger for releasing the restraint of S6_cyto_ from U-motif, allowing rotation and outward movement of S6_cyto_ (Figures 4C and 4G). Two hydrogen bonds (S4167-E4198 and E4198-K4593) newly formed upon opening, seems to be the key for this shift. Other important factors seem to be π-π bond formed by F4171-F4191 (Figures 4G and S5D) and two consecutive glycines (G4179 and G4180) located on the loop connecting the N-terminal and the C-terminal helix of the U-motif (Figure 4G). The former may function as a guide for the shift of the N-terminal helix (Figure 4G inset), and the latter may function as the flexible hinge to enable the deformation of the U-motif.

All alanine-substituted mutants of residues interacting between U-motif and S6_cyto_ as well as two CPVT mutations (N4177S and N4177Y) led to gain-of-function (Figures 4H–4K) with mild enhancement in Ca^2+^ sensitivity and severe loss of Ca^2+^ inactivation (Figures S5F, S5H and S5I). Alanine substitution and pathogenic mutation of F4888 exhibited severe gain-of-function with increased Ca^2+^ sensitivity and loss of Ca^2+^ inactivation (Figures 4L–4O, S5G-S5I). Alanine substitution of F4171, I4172, V4175, and L4914—involved in the hydrophobic pocket—also led to milder gain-of-function of the channel than F4888 mutants (Figures 4L–4O, S5G-S5I). Notably, no additive effects were observed with N4177A and F4888A, indicating that U-motif/S6_cyto_ and U-motif/CTD interactions are involved in the common pathways (Figure S5B). All the residues thought to be involved in the U-motif/S6cyto/CTD interactions and surrounding sequences are conserved in mammalian three RyR subtypes (Figure 4P). These findings suggest that interactions within the U-motif/S6_cyto_/CTD complex play a key role in stabilizing the closed state.

Alanine-substitutions of F4171 and F4191 showed gain-of-function of the channel (Fig. 4M, 4O and S5G-I). No additive effects were observed with double mutant, F4171A_F4191A (Figure S5G), supporting π-π interaction of the two residues. The gain-of-function, however, was opposite to the effect of alanine substitution of S4167, E4198 and K4593, residues involving the compactization of U-motif (Figure 2). This can be explained as follows. In WT, the aromatic rings of F4171 and F4191 form a stable π-π interaction, behaving as a spring which suppresses compaction of the U-motif (Figure S5E). Alanine substitution of F4171 or F4191 weakens the π-π interaction to break the spring, which accelerates compaction of U-motif.

### Structural basis of the loss-of-function mutation

To prove our hypothesis of gating mechanism of the RyR2 channel (Fig. 1H), we conducted structural analysis of a loss-of-function mutant (K4593A). K4593 is the key residue for U-motif-S2-S3 domain interaction (Figure 2A-2C) and compaction of U-motif (Figures 4E-4G). We expected that substitution of this residue with alanine may prevent transduction of the Ca^2+^-binding signal from U-motif to the S2-S3 domain. We have determined the structures of K4593A mutant in the presence of 1 mM EGTA (K4593A(EGTA)) at 3.3 Å resolution (Figure S6A) and in the presence of 100 μM Ca^2+^ (K4593A(Ca^2+^)) at 3.8 Å resolution (Figure S6B). The density for side chain of K4593 completely disappeared in both structures (Figure S6C). The density corresponding to Ca^2+^ was clearly visible at the Ca^2+^ binding site only in the structure of K4593A(Ca^2+^) (Figure S6D). K4593A(EGTA) showed closed state which is very similar to that of WT (Figure 5A and S6E). In contrast, K4593A(Ca^2+^) exhibited large conformational changes around the cytoplasmic region like open state of WT but essentially no change in the TM region (Figure 5B and S6F).

**Fig. 5.**
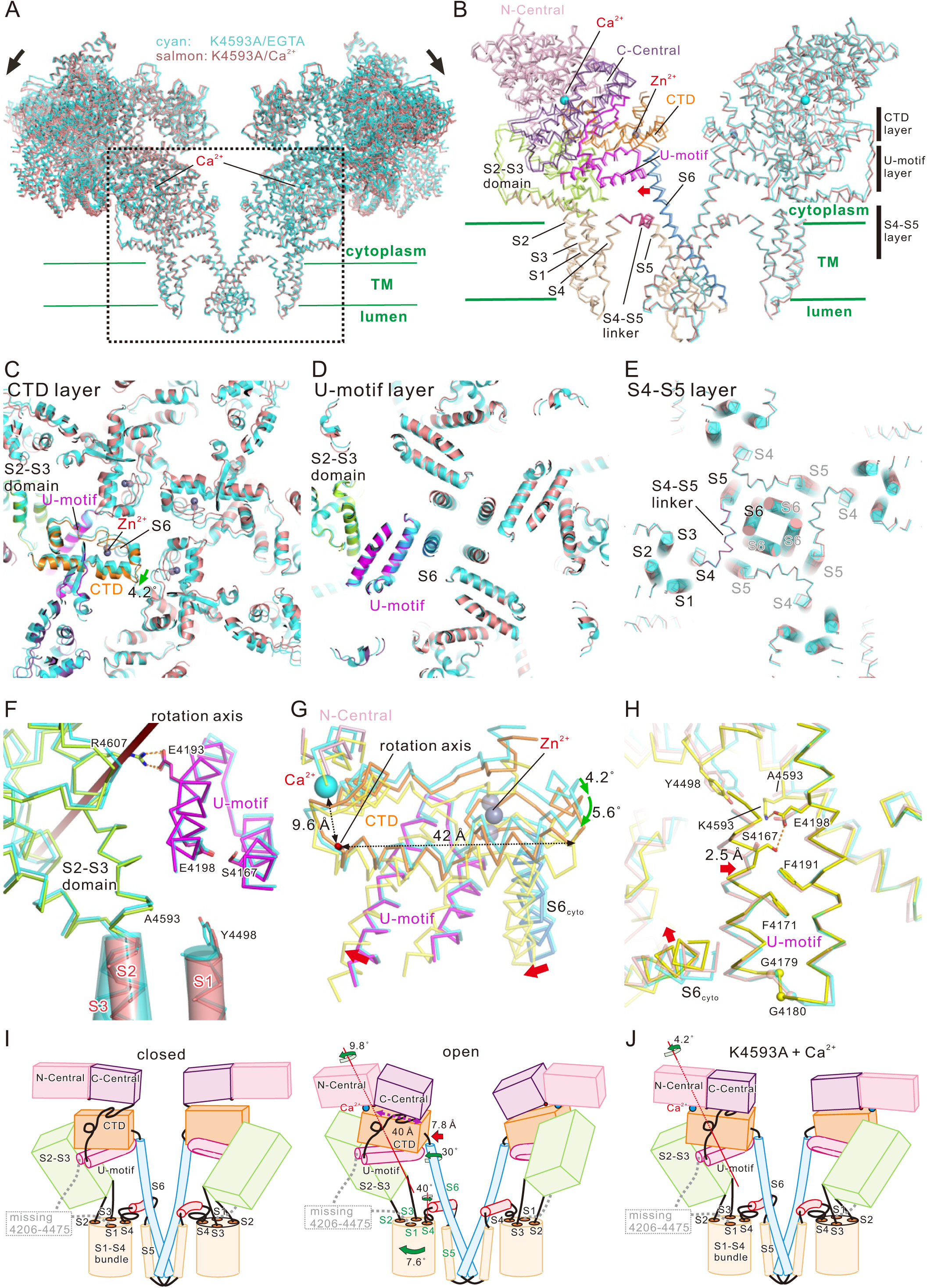
Structural basis of loss-of-function mutation and gating mechanism upon Ca^2+^ binding. (A) Overlay of K4593A mutant in the presence of EGTA (cyan) and K4593A mutant in the presence of Ca^2+^ (salmon) viewed from the direction parallel to the lipid bilayer is shown as a ribbon model. Two facing protomers in the RyR2 tetramer are shown. (B) Magnified view of the dotted box in (A). In the left protomer, each domain is colored (N-Central, light pink; C-Central, purple; U-motif, magenta; S1-S5, wheat; S2-S3 domain, light green; S4-S5 linker, warm pink; S6, blue; CTD, orange). S6 moved outside upon Ca^2+^ binding as indicated by the red arrow. Three regions parallel to the membrane are defined as CTD, U-motif, and S4-S5 layers. Ca^2+^, shown as cyan ball; Zn^2+^, shown as gray ball. (C–E) Cross-section views of CTD, U-motif, and S4-S5 layers. K4593A mutant in the presence of EGTA is colored with cyan and in the presence of Ca^2+^ is colored according to (B) or salmon. In (E), Cα representation overlaid with cylindrical TM helices are used. Ca^2+^ binding causes clockwise rotation of CTD (green arrow in C), but U-motif and S1-S4 TM helices show no considerable movement as shown in Figures 1D and 1E. (F) An overlay of interface of U-motif and S2-S3 domain in K4593A mutant in the presence of EGTA and Ca^2+^ viewed parallel to the membrane and shown as a Cα model. Amino acid residues involved in key interactions are shown as stick models. The color of carbon atoms is the same as that of Cα; oxygen, red; nitrogen, blue. The TM region forming α-helices is overlaid with the cylinder model. Hydrogen bonds or salt bridges are shown as orange dotted lines. (G) Rotation of the C-Central/U-motif/S6_cyto_/CTD complex upon Ca^2+^ binding viewed from the rotation axis. Each domain of K4593A mutant in the presence of Ca^2+^ is colored (U-motif, magenta; S6, blue; CTD, orange). K4593A mutant in the presence of EGTA and WT in the open state is shown as cyan and yellow, respectively. (H) Overlay of three structures (K4593A mutant in the presence of EGTA, K4593A mutant in the presence of Ca^2+^, and WT in the open state) fitted in the C-terminal side of U-motif (4183-4205). K4593A mutant in the presence of EGTA, K4593A mutant in the presence of Ca^2+^, and WT in the open state is shown as cyan, salmon, and yellow, respectively. The N-terminus side of the U-motif of the WT in the open state is ∼2.5 Å closer to the C-terminus side of the U-motif as indicated by the red arrow, and as a result, S6_cyto_ movement (red arrow) becomes possible. This movement was not observed in the K4593A mutant in the presence of Ca^2+^. (I, J) Scheme of the structure in the closed (left), open state (right) (I) and K4593A mutant in the presence of Ca^2+^ (J). In the closed state, the upper part of S4 does not form an α-helix. The S4-S5 linker is unfolded and significantly bends in the direction of S6. In the open state, binding of Ca^2+^ to the C-Central/CTD interface causes 9.8° rotation of all domains consisting of C-Central/U-motif/S6_cyto_/CTD complex and compactization of U-motif, leading to two pathways. Pathway 1: the rotation causes 30º rotation of S6_cyto_ which loosens the U-motif/S6_cyto_ interaction and allows outward movement of S6. Pathway 2: a sequential movement of the S2S3 domain, S2, S1-S4 bundle, and S4 allows the upper part of S4 to rewind and form an α-helix. Subsequently, the S4-S5 linker moves outward, creating a space where S6 can lean into. A combination of these two independent pathways eventually leads to opening of the channel. (J) In the K4593A mutant, binding of Ca^2+^ to the C-Central/CTD interface causes 4.2° rotation of the C-Central/U-motif complex, but no considerable movement occurs in the U-motif and S6_cyto_, therefore, the signal of Ca^2+^ binding is not transmitted to the subsequent pathway, and the opening of the channel pore does not occur.

Changes associated with Ca^2+^ binding were analyzed at three layers (Figures 5C-E; Video S6). Upon Ca^2+^ binding, CTD rotated clockwise (4.2°, Figure 5C). In contrast, U-motif and the TM helices hardly moved (Figures 5D-5E). Although a slight movement was observed in the upper part of S2-S3 domain in the structure of K4593A(Ca^2+^) compared to the structure in K4593A(EGTA) (Figure 5F), essentially no movement was detected juxtamembrane region of S1-S4 bundle (Figure 5F). The rotation of CTD in K4593(Ca^2+^) (4.2°) is about half of that in WT in the open state (9.8°, Figure 5G). No compaction of U-motif was detected in K4593A(Ca^2+^) (Figures 5H), indicating that the hydrogen bond between K4593 and E4198 is indispensable for the compaction of U-motif. The subsequent outward movement of S6_cyto_ was not detected either (Figures 5H). On the other hand, no considerable difference was detected between K4593A(EGTA) and WT in the closed state at three layers (Figure S6G) nor the shape of U-motif (Figure S6H), supporting that the above changes detected in K4593A(Ca^2+^) was derived from the effect of K4593A mutation. These lines of evidence support our hypothesis that compaction and rotation of U-motif move S2-S3 domain to open the channel pore.

### Summary of the Ca^2+^ induced movements in the RyR2

Here, we summarize the channel opening mechanism associated with Ca^2+^ binding of RyR2 determined by high-resolution cryo-EM structures: (1) binding of Ca^2+^ rotates the U-motif/S6_cyto_/CTD complex by 9.8° along with the axis which is 40 Å apart from S6_cyto_; (2) compaction of U-motif; (3) extruding the S2-S3 domain by the U-motif causes S2 to move; (4) movement of S2 rotates the S1-S4 bundle by 7.6° to release the stopper comprising F4749/L4505; (5) rotation of the upper part of S4 allows outward movement of the S4-S5 linker, creating a space where S6 can lean into; and (6) rotation of S6_cyto_ by 30° causes the outward leaning of S6 by 7.8 Å (Figures 5I). Thus, the sequential rotations of domains/α-helices and domain-domain interactions may occur in conformation changes associated with Ca^2+^ binding. Rotation of U-motif in K4593A in the presence of Ca^2+^ stopped in the middle of the transition from the closed state to the open state, that is, binding of Ca^2+^ rotates only U-motif by 4.2° along with the same axis as the rotation from closed state to open state (Figure 5J).

## Discussion

By combining high-resolution cryo-EM structures and quantitative functional analysis of mutant channels, we successfully clarified the gating mechanism of RyR2 upon Ca^2+^ binding. Figure 6A shows a schematic diagram of the channel core domain of RyR2 with details of the important interactions identified in this study. Mutations of residues involved in the signaling pathway for outward movement of the S4-S5 linker caused loss-of-function, whereas those supporting the stopper (i.e., S1/S3/S4 and S1/S4 interactions) or the U-motif/CTD/S6_cyto_ complex led to gain-of-function. Given the locations of these mutations, it becomes apparent that interactions close to the channel pore are important for stabilizing the channel in the closed state and those in the surrounding region are essential for channel opening.

**Fig. 6.**
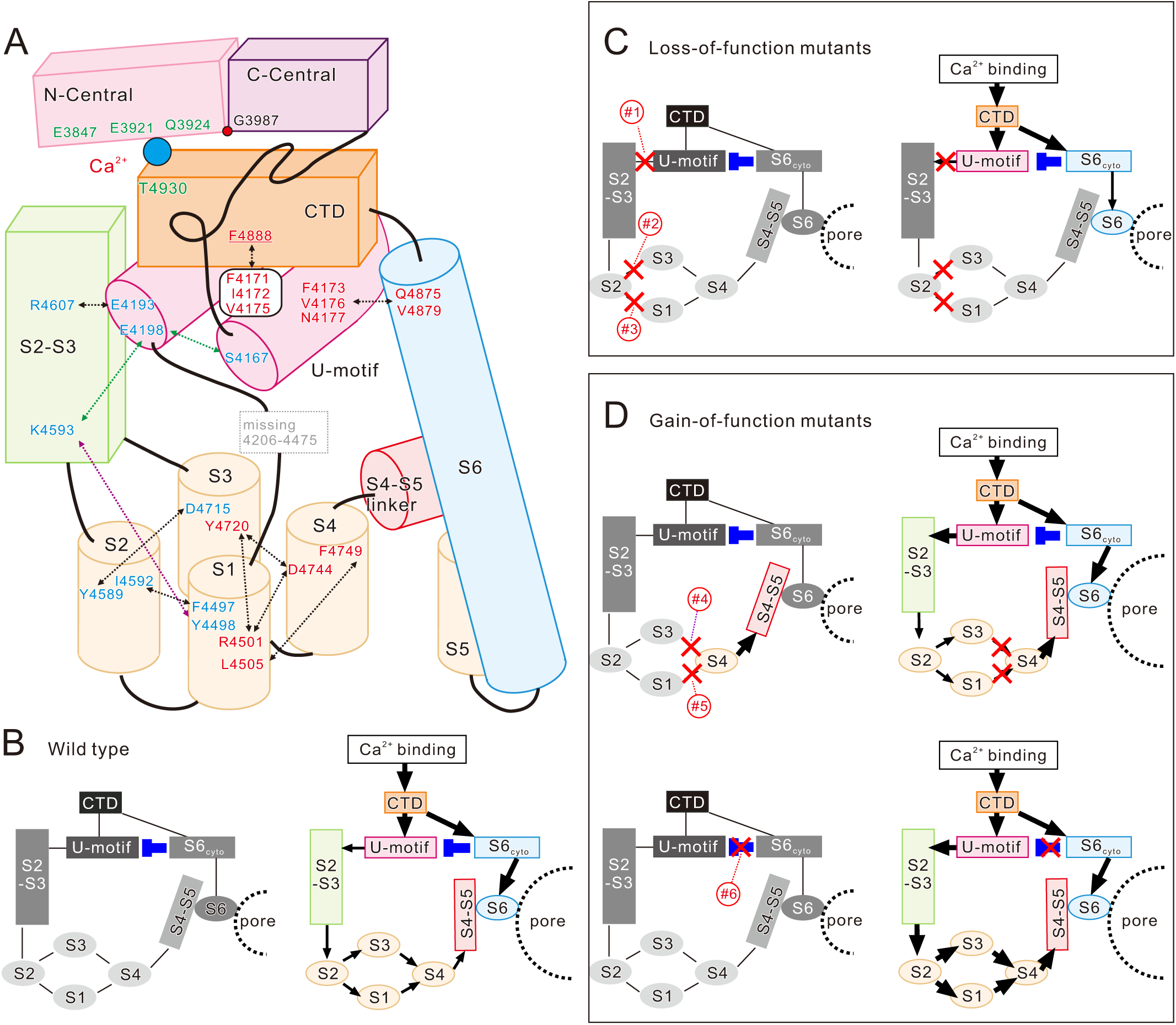
Details of interactions and schematic diagram of the RyR2 channel gating mechanism upon Ca^2+^ binding. (A) Details of interactions identified in this study. Amino-acid residues shown in red letter and blue letter indicate gain-of-function and loss-of-function by alanine-substituted or pathogenic mutations, respectively. Arrows indicate interactions. Yellow and dark blue arrows indicate interactions only found in the closed and open states, respectively. (B-D) Schematic diagram of the RyR2 channel gating mechanism upon Ca^2+^ binding. The left and right diagrams show the states in the absence and presence of Ca^2+^, respectively. The black lines, the domain interactions; The blue T-shaped lines, the domain interactions that act as the suppression; The black lines with arrowhead, the activated domain interactions. The domains shown in gray-scale and colored indicate the domains in the inactive and active states, respectively. (B) Wild type (WT). Since the movement of S4-S5 linker is locked, the channel pore is closed. Ca^2+^ binding unlocks the S4-S5 linker and induces the outward leaning of S6, resulting in the pore opening. (C) Loss-of-function (LOF) mutants. Mutations in the U-motif/S2-S3 domain (#1), S1/S2 (#2) or S2/S3 (#3) interface cause disconnection of signal transduction. Binding of Ca^2+^ therefore cannot induce the outward movement of the S4-S5 linker and the channel pore is kept closed. (D) Gain-of-function (GOF) mutants. (Upper panels) Mutations in S3/S4 (#4) or S1/S4 (#5) interface unlock the S4-S5 linker to be activated. In the absence of Ca^2+^, the channel pore is kept closed, since the outward leaning of S6 does not occur spontaneously. Binding of Ca^2+^ causes hyperactivity of the channel, since the S4-S5 linker is more active than WT. (Lower panels) Mutations in U-motif, S6_cyto_ or CTD (#6) reduce or lose U-motif/S6_cyto_ interaction. Binding of Ca^2+^ causes hyperactivity of the channel, since S6 and the S4-S5 linker is more active than WT.

The present results can reasonably explain the gating mechanism of RyR2 channel and how pathogenic or alanine mutations affect the channel activity (Figures 6B-6D). In the closed state of the WT channel, the S4-S5 linker prevents outward leaning of S6 (Figure 6B, left). In addition, S6_cyto_ restricts movement of the U-motif toward the S2-S3 domain. In the open state of the WT, binding of Ca^2+^ moves CTD which transmits signals into two downstream pathways to open the channel; S4-S5 linker is unlocked by sequential conformational changes beginning at the U-motif and S6 leans outward by the movement of S6_cyto_. However, since S6_cyto_ still restricts the U-motif, the activity may be medium (Figure 6B, right). Mutations in the U-motif/S2-S3 domain, S1/S2, or S2/S3 interface disrupt the interaction which is essential for the channel opening (Figure 6C, left). Upon Ca^2+^ binding, conformational changes stop on the way, resulting in loss-of-function (Figure 6C, right). Mutations in S3/S4 or S1/S4 interface unlock the S4-S5 linker (Figure 6D, upper left). However, in the absence of Ca^2+^, this does not induce spontaneous outward movement of the S4-S5 linker and the channel pore kept closed. In the presence of Ca^2+^, these channels will show gain-of-function, since the S4-S5 linker is more mobile (Figure 6D, upper right). Mutations in the U-motif/S6_cyto_ or the U-motif/CTD disrupt the U-motif/S6_cyto_ interaction to make the U-motif and S6_cyto_ more mobile (Figure 6D, lower left). Upon Ca^2+^ binding, these channels will show gain-of-function by an increased mobility in the S4-S5 linker and S6_cyto_ (Figure 6D, lower right). Considering that residues involved in the interactions are highly conserved and pathogenic mutations in RyR1 (L4505P, D4744H, and F4888Y; see Table 1) also showed corresponding behaviors, the fundamental mechanism of channel gating may be common among three RyR subtypes.

Many structures of pig RyR2 have been determined (Chi et al., 2019; Gong et al., 2019; Peng et al., 2016). Among them, structures in the open state have been classified into two groups: PCB95 in addition to Ca^2+^ (Ca^2+^/PCB95) (Peng et al., 2016) and ATP/caffeine in addition to Ca^2+^ (Ca^2+^/ATP/caffeine) (Chi et al., 2019). We compared our structure in the open state with those structures by analyzing conformational changes upon Ca^2+^ binding at CTD layer and at U-motif layer (Figure S7A). In the Ca^2+^/PCB95 structure, rotation of CTD and U-motif occurred upon opening, thereby outward movement of S6 occurred (Figure S7A 1^st^ row). In the Ca^2+^/ATP/caffeine structure, in contrast, both CTD and U-motif made translational outward movement (Figure S7A 2^nd^ row). Conformational change in our structures was similar to that observed in the Ca^2+^/PCB95 structure (Figure S7A 3^rd^ row). Ca^2+^/PCB95 open structure was only determined in the absence of FKBP12.6; addition of FKBP12.6 closes the channel (Chi et al., 2019). This is contrast to our open structure which binds FKBP12.6 (Figure S1I).

Previous structures of RyR in the presence of Ca^2+^ alone failed to get open state; the channel pore was closed (Chi et al., 2019; des Georges et al., 2016). Instead, slight movements of CTD, U-motif, and S6 occurred upon Ca^2+^ binding (referred to as ‘primed’ state). Analysis of conformational changes in CTD and U-motif layers revealed slight rotation of CTD and U-motif and slight outward movement of S6 in the primed state, indicating an intermediate state between closed and open states (Figure S7B 1^st^ row). In contrast, our K4593A(Ca^2+^) structure showed no movement of U-motif and S6 at all, reflecting disruption of transduction signal from Ca^2+^ binding (Figure S7B 2^nd^ row). Thus, K4593A(Ca^2+^) is an artificial state caused by specific mutation and different from the reported primed state (Hirose et al., 2021).

Outward movement of the S4-S5 linker is important in the channel opening, as it creates the space where S6 can lean into (des Georges et al., 2016; Peng et al., 2016). des Georges et al. (des Georges et al., 2016) provided a more detailed description about the movement of the S4-S5 linker in RyR1; it is bent in the direction of S6 in the closed state, but it straightens and alters its contacts with S6, thereby opening the channel pore. We showed similar conformational changes in the S4-S5 linker in RyR2 and further demonstrated that rewinding of the upper part of S4 to an α-helix, which is caused by rotation of the S1-S4 bundle, drives the S4-S5 linker movement (Figure 3H-3J and S4D-S4F). Regulatory mechanisms of interaction between the S4-S5 linker and S6 will be the next subject for complete understanding of channel gating.

In RyR1, rearrangement of the salt bridges between S1 (R4563), S3 (Y4791) and S4 (D4815) has been reported to occur upon channel opening (des Georges et al., 2016). We showed that the corresponding salt bridges between R4501, Y4720, and D4744 were preserved in RyR2 but maintained in both closed and open states (Figure 3A-3C and S4A-S4C). Functional assays revealed that hydrogen bonds/salt bridges are critical to keep the stopper in place, as disruption of salt bridges by alanine substitutions or pathogenic mutations caused severe gain-of-function of the channel (Figure 3D-3G and S4G). Interestingly, a recent study showed that diamide insecticides bind to the pocket within the S1-S4 bundle of RyR1, thereby disrupting the salt bridge and opening RyR1 (Ma et al., 2020). Our results reasonably explain why such the insecticides activate the RyR channel.

The U-motif is proposed to be important in channel gating as it forms a stable complex with S6_cyto_ and CTD, which move together when the channel opens (Yan et al., 2015). This movement is accompanied by a rigid shift in the S2-S3 domain, which has been proposed to eventually open the channel pore (Bai et al., 2016). We found that interaction of the U-motif with the S2-S3 domain is critically important in the channel opening; disruption of the interaction causes loss-of-function of the channel (Figure 2 and S3). In addition, we demonstrated that the U-motif/S6_cyto_ interaction stabilizes the channel to the closed state (Figure 4 and S5). Stabilization of the channel is important not only for prevention of the hyperactivity but also for regulation by various stimuli or modifications that activate the channel, e.g., phosphorylation or oxidation/S-nitrosylation (Benkusky et al., 2004; Santulli et al., 2018). The U-motif/S6_cyto_ interaction is the stabilization mechanism first identified at the molecular level and thus may contribute to regulation of the RyR channel.

In RyRs, the large cytoplasmic region is believed to play an important role in regulating channel opening. Indeed, a number of pathogenic mutations is located in the cytoplasmic region. The question arises as to how these mutations, located far from the core domain, cause changes in the channel activity. We demonstrated that the C-Central domain forms a tight complex with the U-motif/S6_cyto_/CTD to move together upon Ca^2+^ binding (Figures 1F, S2E, and S2F). The N-Central domain also moves upon Ca^2+^ binding (Figures 1A and 1F). Through the N-/C-Central domains, the core domain has a series of interactions with the N-terminal domains (NTD), HANDLE, and HD1 domains which consist a major part of the large cytoplasmic region (Peng et al., 2016; Yan et al., 2015). Thus, it is possible that the conformational changes of the cytoplasmic region by pathogenic mutations affect the channel gating through these interactions. Recently, RyR1 and RyR2 structures carrying gain-of-function mutations at N-terminal domains (NTD) were reported (Iyer et al., 2020). Determining the structures of various mutant RyRs will help to elucidate the long-range allosteric gating mechanism in the future.

In conclusion, we have revealed the gating mechanism of RyR2 upon Ca^2+^ binding and provided structural insights into the effects of pathogenic mutations on channel activity. These findings may greatly help develop more effective drugs to treat RyR2-associated diseases.

## Acknowledgements

We thank Mariko Kurakata for assistance with cell culture. We thank Ikue Hiraga and the Laboratory of Radioisotope Research, Research Support Center, Juntendo University Graduate School of Medicine, for technical assistance. We also thank the staff scientists at the University of Tokyo’s cryo-EM facility. This study was partly supported by the Japan Society for the Promotion of Sciences KAKENHI (grant number JP16H04748 to H.O., 19K07105 to N.K., and 19H03404 to T.M.); the Platform Project for Supporting Drug Discovery and Life Science Research (Basis for Supporting Innovative Drug Discovery and Life Science Research [BINDS]; grant number JP20am0101080 to H.O. and T.M.); the Practical Research Project for Rare/Intractable Diseases from the Japan Agency for Medical Research and Development (AMED; grant number 19ek0109202) to N.K.); an Intramural Research Grant (29-4 and 2-5) for Neurological and Psychiatric Disorders of NCNP (to T.M.); and the Vehicle Racing Commemorative Foundation (to H.O. and T.M.).

## Author contributions

T.K., N.K., T.M., and H.O. conceived and designed the project. M.Ko. and H.O. performed cell culture. T.K. performed protein purification. K.S. performed negative-staining EM studies. A.T. and T.K. prepared the grid for cryo-EM. A.T. and M.Ki. processed the images. H.O. performed model building and refinement. T.M. and N.K. performed the functional analysis. T.K., A.T., N.K., K.S., M.Ko., T.S. M.Ki., T.M., and H.O interpreted the data. T.M. and H.O supervised the project. H.O. and T.M. wrote the manuscript with input from all authors.

## Declaration of interests

The authors declare no competing interests.

## Materials and methods

### Expression and purification of SBP-FKBP12.6

Expression and purification of SBP-tagged FKBP12.6 was performed as described previously (Cabra et al., 2016) with some modifications. Briefly, cDNA for SBP-tagged human FKBP12.6 was subcloned into the pET28 vector (Novagen) containing a 6×His-tag at the N-terminus. *Escherichia coli* BL21 cells transformed with the above expression vector were grown at 37°C in 500 mL LB medium containing ampicillin at a final concentration of 50 mg/L. After reaching an OD_600_ of 0.95–1.2, SBP-FKBP12.6 expression was induced by adding 1 mM IPTG (Wako) for 3 h at 37°C. Cells were then harvested, resuspended in 20 mM MOPS-Na (pH 7.4), 300 mM NaCl, 20 mM imidazole, and a cocktail of protease inhibitors [antipain (Peptide Institute), aprotinin (Nacalai), chymostatin (Peptide Institute), leupeptin (Peptide Institute) and pepstatin A (Peptide Institute), each 2 μg/mL], and lysed using a sonicator in an ice bath. The lysate was centrifuged at 100,000 × *g* for 30 min at 4 °C and the pellet was discarded. The supernatant was incubated for 1 h at 4°C with 0.5 mL Profinity IMAC resin (Bio-Rad Laboratories, Hercules, CA) in a buffer containing 20 mM MOPS-Na (pH 7.4), 300 mM NaCl, and 20 mM imidazole. The resin was washed five times with the buffer and SBP-FKBP12.6 was eluted with buffer containing 300 mM imidazole. The eluted protein was quickly frozen in liquid nitrogen and stored at −80 °C.

### Construction of mutant RyR2 cDNA expression vector and generation of stable HEK293 cell lines

Alanine substitution or pathogenic mutations were introduced by inverse PCR using F5 cassette (nucleotide residue number 10,185-14,901 of mouse RyR2) as a template (Uehara et al., 2017). The mutations were confirmed by DNA sequencing. The mutant cassette was inserted into full-length wild-type mouse RyR2 expression vector (pcDNA5/FRT/TO-RyR2wt) (Uehara et al., 2017) by restriction enzymes (*BsiW* I/*Not* I). HEK293 cells stably expressing mutant RyR2 was generated using the Flp-In T-REx system (ThemoFisher) according to manufacturer’s instructions. Clones with suitable expression of RyR2 were selected and used for experiments.

### Preparation of microsomes from RyR2-expressing HEK293 cells

HEK293 cells stably expressing mouse RyR2 (wild type or mutants) were grown in 60 150-mm cell culture dishes. At 70–80% confluency, protein expression was induced by 2 μg/mL doxycycline (SIGMA) for 48 h. Cells were then harvested, rinsed with cold phosphate-buffered saline (PBS) (GIBCO), and microsomes were prepared as described by Inesi et al. (Eletr and Inesi, 1972). Briefly, the cell pellet was resuspended in 60 mL of 10 mM NaHCO_3_ in presence of protease inhibitors and processed for nitrogen cavitation for 30 min at 1,000 psi. The suspension was diluted with 60 mL of 0.6 M sucrose, 0.3 M KCl, 40 mM MOPS-Na (pH 7.4), and protease inhibitors and then centrifuged at 1,000 × *g* for 10 min. The supernatant was supplemented with 30 mL of 2.4 M KCl, 0.3 M sucrose, 20 mM MOPS-Na (pH 7.4), and the above protease inhibitor cocktail and centrifuged at 10,000 × *g* for 20 min. The supernatant was then ultracentrifuged at 100,000 × *g* for 30 min. The microsomal pellet was resuspended in 60 mL of 0.6 M KCl, 0.3 M sucrose, 20 mM MOPS-Na (pH 7.4), and the protease inhibitor cocktail, and ultracentrifuged again. Finally, the pellet was resuspended in 12 mL of 0.3 M sucrose, 20 mM MOPS-Na (pH 7.4), and the protease inhibitor cocktail, followed by quick freezing in liquid nitrogen and storage at −80 °C until further use.

### Purification of RyR2

RyR2 was purified using SBP-FKBP12.6 affinity chromatography (Cabra et al., 2016) with some modifications. The microsomes were solubilized with 2% (w/v) CHAPS (Dojindo) and 1% (w/v) soybean lecithin (Avanti Polar Lipids) in 1 M NaCl, 20 mM MOPS (pH 7.4), 2 mM dithiothreitol, and the protease inhibitor cocktail for 30 min on ice. After centrifugation at 100,000 × *g* for 30 min at 4 °C, the supernatant was diluted with four volumes of 20 mM MOPS (pH 7.4), 2 mM dithiothreitol, and the protease inhibitor cocktail, after which it was passed through a 0.45-μm filter and loaded onto a pre-equilibrated 5-mL StrepTrap HP column (GE Healthcare, Chicago, IL) with bound SBP-FKBP12.6 fusion protein. The column was successively washed with 10 column volumes of (1) wash buffer (20 mM MOPS pH7.4, 2 mM dithiothreitol, and 0.3 M sucrose) containing 0.2 M NaCl and 0.25% (w/v) CHAPS and (2) wash buffer containing 0.5 M NaCl and 0.015% (w/v) Tween-20 (Sigma-Aldrich). The SBP-FKBP12.6-RyR2 complex was eluted with the wash buffer supplemented with 2.5 mM D-desthiobiotin (Iba Lifesciences). After checking the purity by SDS-PAGE, the eluate was quickly frozen in liquid nitrogen and stored at −80 °C until further use.

### Negative staining

The purified RyR2 sample was diluted with a buffer containing 0.2 M NaCl, 20 mM MOPS-Na (pH 7.4), 2 mM dithiothreitol, and 0.015% (w/v) Tween-20, and then applied to pre-hydrophilized carbon-coated EM grids (400 mesh hexagonal copper grids, Stork Veco BV, Netherlands), negatively stained with 1.4% (w/v) uranyl acetate solution, and observed at 40,000× magnification using a transmission electron microscope (H7500; Hitachi High-Technologies, Tokyo, Japan) operating at 80 kV. Micrographs were taken at 40,000× using a 1,024 × 1,024 pixel CCD camera (Fast Scan-F114; TVIPS, Gauting, Germany).

### Cryo-EM sample preparation

The purified RyR2 sample was buffer-exchanged and concentrated to ∼5 mg/mL using Amicon Ultra 100k (Millipore, Burlington, MA) with a buffer containing 0.5 M NaCl, 20 mM MOPS-Na (pH 7.4), 2 mM dithiothreitol, and 0.015% (w/v) Tween-20. EGTA (final concentration of 1 mM) or CaCl_2_ (final concentration of 100 μM) was added to the concentrated protein samples to fix the channel to the closed or open state. The concentrated RyR2 was loaded onto a Quantifoil Cu/Rh grid (R1.2/1.3, 300 mesh) (Quantifoil), blotted using Vitrobot Mark III (FEI) with 4 s of blotting time and 100% humidity at 6 °C, and then plunge-frozen in liquid ethane.

### Cryo-EM data collection

Grids were screened for ice quality and cryo-EM data were acquired using a Titan Krios G3i cryo-EM (Thermo Fisher Scientific, Waltham, MA) running at 300 kV and equipped with a Gatan Quantum-LS Energy Filter (slit width 25 eV) and a Gatan K3 Summit direct electron detector in the electron counting mode. The electron flux was set to 14 e^-^/pix/s at the detector. For the WT close state with EGTA and F4888A with EGTA, imaging was performed at a nominal magnification of 87,000×, corresponding to a calibrated pixel size of 1.07 Å/pix (University of Tokyo, Japan). Electron dose was set to 50 e^-^/Å^2^ for the WT close state and 48 e^-^/Å^2^ for F4888A with EGTA. For other samples, imaging was performed at a nominal magnification of 105,000×, corresponding to a calibrated pixel size of 0.83 Å/pix. Electron dose was set to 60 e^-^/Å^2^. Each movie was subdivided into 1 e^-^/frame for all datasets. The data were automatically acquired by the image shift method using SerialEM software (Mastronarde, 2005), with a defocus range of -0.8 to -1.6 μm.

### Image processing

Unless otherwise stated, the same procedure was used to process the data. The movie frames were motion corrected by RELION and CTF parameters and estimated by CTFFIND4 (Rohou and Grigorieff, 2015). All the following processes were performed using RELION (ver. 3.0 and 3.1) (Zivanov et al., 2018). To generate 2D templates for automatic particle picking, particles were picked from 100 randomly selected micrographs using template-free Laplacian-of-Gaussian picking, then subjected to multiple rounds of reference-free 2D classification. Good 2D classes were selected as templates and 2D template-based particle picking was performed.

Picked particles were extracted with 4× down-sampling and subjected to one round of 2D classification. Selected good particles from the 2D classification were submitted to 3D classification. After one round of 3D classification, good particles were selected, re-extracted with 1.5× down-sampling, and subjected to 3D refinement. The resulting 3D map and particle set were then subjected to beam-tilt and per-particle defocus refinement, Bayesian polishing, and second round of per-particle defocus refinement followed by 3D refinement. To separate the particles based on open/close states, the no-align 3D classification was performed using a mask covering the TM region. Good classes were subjected to final 3D refinement and postprocessing. Resolution was determined according to the Fourier shell collation (FSC) = 0.143 criterion.

For F4888A EGTA, two datasets were acquired, and particles were combined after Bayesian polishing. In each dataset, particles were selected from one round of 3D classification following two rounds of 2D classification. Detailed information is listed in Supplementary table 1 and described in figs. S2, S13, and S14.

### Modelbuilding, refinement, and analysis

Model building was performed using COOT (Emsley et al., 2010). RyR1 in the closed state (PDB accession code, 5TB0) was used as the initial reference model. The coordinates were rigid-body fitted in UCSF Chimera (Pettersen et al., 2004). After substitutions in mouse RyR2 sequence and manual building of the model, real space refinement was performed with PHENIX (Adams et al., 2010; Afonine et al., 2018) with secondary structure and geometry restrained. The residue sequences (1– 10, 85–108, 861–864, 954–969, 1,015–1,026, 1,063–1,083, 1,275–1,283, 1,447–1,565, 1,851–1,890, 2,010–2,055, 2,362–2,378, 2,443–2,451, 2,659–2,711, 2,757–2,760, 2,785–2,834, 2,906–2,915, 2,943–2,960, 3,030–3,103, 3,130–3,135, 3,221–3,230, 3,435–3,476, 3,580–3,610, 3,649–3,658, 3,700–3,711, 4,206–4,272, 4,312–4,477, 4,522–4,555, and 4,963–4,966) were omitted, as the corresponding densities were not visible in all of the maps. All figures were prepared using PyMOL (The PyMOL Molecular Graphics System; http://www.pymol.org). Pore radii along the ion conducting pathway were calculated with HOLE (Smart et al., 1993). Buried surface areas were calculated with CNS (Brünger et al., 1998).

### [^3^H]Ryanodine binding

[^3^H]Ryanodine binding assay was carried out as described previously (Fujii et al., 2017; Murayama et al., 2018a). Briefly, microsomes prepared from HEK293 cells expressing RyR2 were incubated for 1 h at 25 °C with 5 nM [^3^H]ryanodine (PerkinElmer) in reaction media containing 0.17 M NaCl, 20 mM MOPSO-Na (pH 7.0), 2 mM dithiothreitol, 1 mM AMP, and 1 mM MgCl_2_. Free Ca^2+^ was adjusted with 10 mM EGTA using WEBMAXC STANDARD (https://somapp.ucdmc.ucdavis.edu/pharmacology/bers/maxchelator/webmaxc/webmaxcS.htm). The [^3^H]ryanodine binding data (*B*) were normalized to the maximum number of functional channels (*B*_max_), which was separately determined by Scatchard plot analysis using various concentrations (3– 20 nM) of [^3^H]ryanodine in a high-salt medium. The resultant *B*/*B*_max_ represents the averaged activity of each mutant.

### Single-cell Ca^2+^ imaging

Single-cell Ca^2+^ imaging was performed using HEK293 cells expressing WT or mutant RyR2. In Ca^2+^ measurements with RyR2, the Ca^2+^ signals from the cytoplasm ([Ca^2+^]_cyt_) and ER lumen ([Ca^2+^]_ER_) were monitored using G-GECO1.1 (a gift from Robert Campbell from University of Alberta; Addgene plasmid #32445) (Zhao et al., 2011). and R-CEPIA1er (a gift from Masamitsu Iino, Nihon University, Tokyo, Japan; Addgene plasmid #58216) (Suzuki et al., 2014), respectively. Cells were transfected with cDNAs for these Ca^2+^ indicators 26–30 h before measurement and at the same time doxycycline (2 μg/mL) was added to the culture medium to induce RyR2 expression. Experiments were carried out in HEPES-buffered Krebs solution (140 mM NaCl, 5 mM KCl, 2 mM CaCl_2_, 1 mM MgCl_2_, 11 mM glucose, and 5 mM HEPES at pH 7.4). At the end of each experiment, *F*_min_ and *F*_max_ were obtained with 0Ca Krebs solution containing 20 μM ionomycin, 5 mM BAPTA, and 20 μM cyclopiazonic acid and 20Ca Krebs solution containing 20 μM ionomycin and 20 mM CaCl_2_, respectively (Uehara et al., 2017). The fluorescence signal (*F*−*F*_min_) was normalized to the maximal fluorescence intensity (*F*_max_−*F*_min_). Measurements were carried out at 26 °C.

### Titles for multimedia files

**Video S1**.

The architecture of RyR2 and details of its opening.

**Video S2**.

Rotation of C-Central/U-motif/U-motif/S6_cyto_/CTD complex upon Ca^2+^ binding.

**Video S3**.

Details of signal transmission from the cytoplasm into the TM region via the U-motif/S2-S3 domain interaction.

**Video S4**.

Conformation changes in the TM region upon Ca^2+^ binding.

**Video S5**.

Details of the U-motif/S6_cyto_/CTD interaction.

**Video S6**.

Structure comparison of the K4593A mutant with the WT in the closed state and open state.

## Notes

### Competing Interest Statement

The authors have declared no competing interest.

### Summary of Updates

Some modifications of main text to clarify; Figures 2, 3, 4, 5, 6 revised; Supplemental files revised

