## Supplemental Figures and table for "Molecular basis for gating of cardiac ryanodine receptor: underlying mechanisms for gain- and loss-of function mutations"

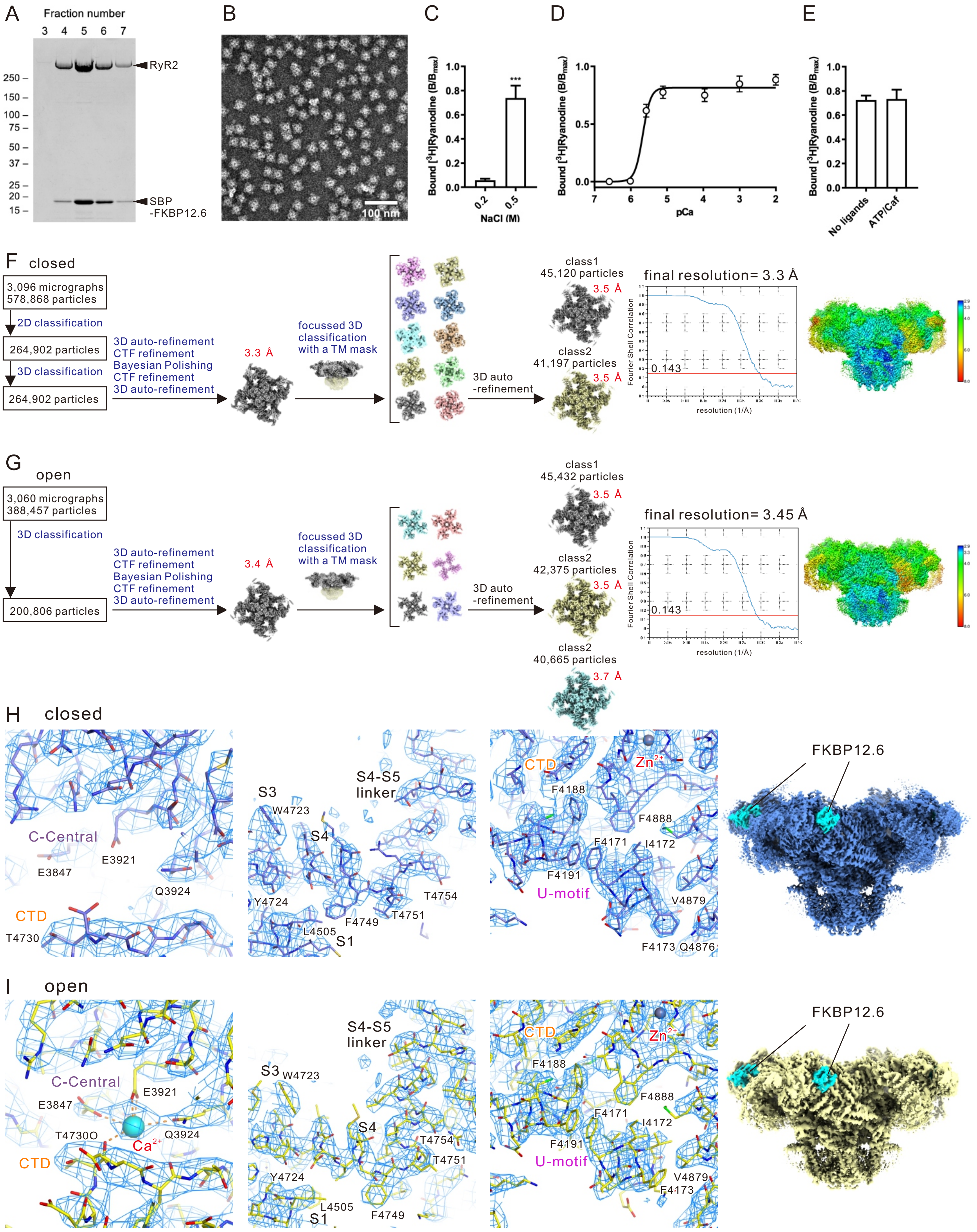
A
Fraction number
3 4 5 6 7
RyR2
SBP -FKBP12.6
250 150 100 75 50 37 25 20 15
B
100 nm
C
Bound [3H]Ryanodine (B/Bmax)
***
NaCl (M)
D
pCa
E
No ligands
ATP/Caf
F closed
3,096 micrographs
578,868 particles
2D classification
264,902 particles
3D classification
264,902 particles
3D auto-refinement
CTF refinement
Bayesian Polishing
CTF refinement
3D auto-refinement
3.3 Å
focussed 3D classification with a TM mask
3D auto -refinement
class1
45,120 particles
3.5 Å
class2
41,197 particles
3.5 Å
final resolution= 3.3 Å
Fourier Shell Correlation
0.143
resolution (1/Å)
G open
3,060 micrographs
388,457 particles
3D classification
200,806 particles
3.4 Å
class1
45,432 particles
class2
42,375 particles
class2
40,665 particles
3.7 Å
final resolution= 3.45 Å
H closed
C-Central
E3847
E3921
Q3924
CTD
T4730
S3
W4723
S4-S5 linker
S4
Y4724
T4754
T4751
F4749
L4505
S1
Zn2+
F4188
F4888
F4171
I4172
F4191
U-motif
V4879
F4173 Q4876
FKBP12.6
I open
T4730O
Ca2+

Figure S1

**Figure. S1. Single particle analysis on the closed and open state, related to Figure 1.**

(A) SDS-PAGE of the peak fractions of StrepTrap column chromatography.

(B) Representative image of the purified RyR2 complex by negative staining.

(C) Effect of NaCl concentration on [$^3$H]ryanodine binding of the purified RyR2. Data are presented as the mean ± SD (n = 6). ***p < 0.001.

(D) $Ca^{2+}$ dependence on [$^3$H]ryanodine binding of the purified RyR2 in the presence of 100 μM $Ca^{2+}$. Data are presented as the mean ± SD (n = 6).

(E) Effect of ATP and caffeine on [$^3$H]ryanodine binding of the purified RyR2. Data are presented as the mean ± SD (n = 3).

(F and G) Workflows for cryo-EM data processing and estimations of resolution by Fourier shell correlation (FSC) plots and local resolution EM maps in the presence of EGTA (F; closed state) and in the presence of 100 μM $Ca^{2+}$ (G; open state), respectively.

(H and I) Density maps for the reconstructed structures in the presence of EGTA (H; closed state) and in the presence of 100 μM $Ca^{2+}$ (I; open state), respectively. From left to right: around the $Ca^{2+}$ binding site, contour level at 0.03: around the S4-S5 linker, contour level at 0.035: around the U-motif, contour levels for the closed and open states were 0.035 and 0.03, respectively: overall density and the density corresponding to FKBP12.6 is colored with cyan. contour levels for the closed and open states were 0.023 and 0.017, respectively.

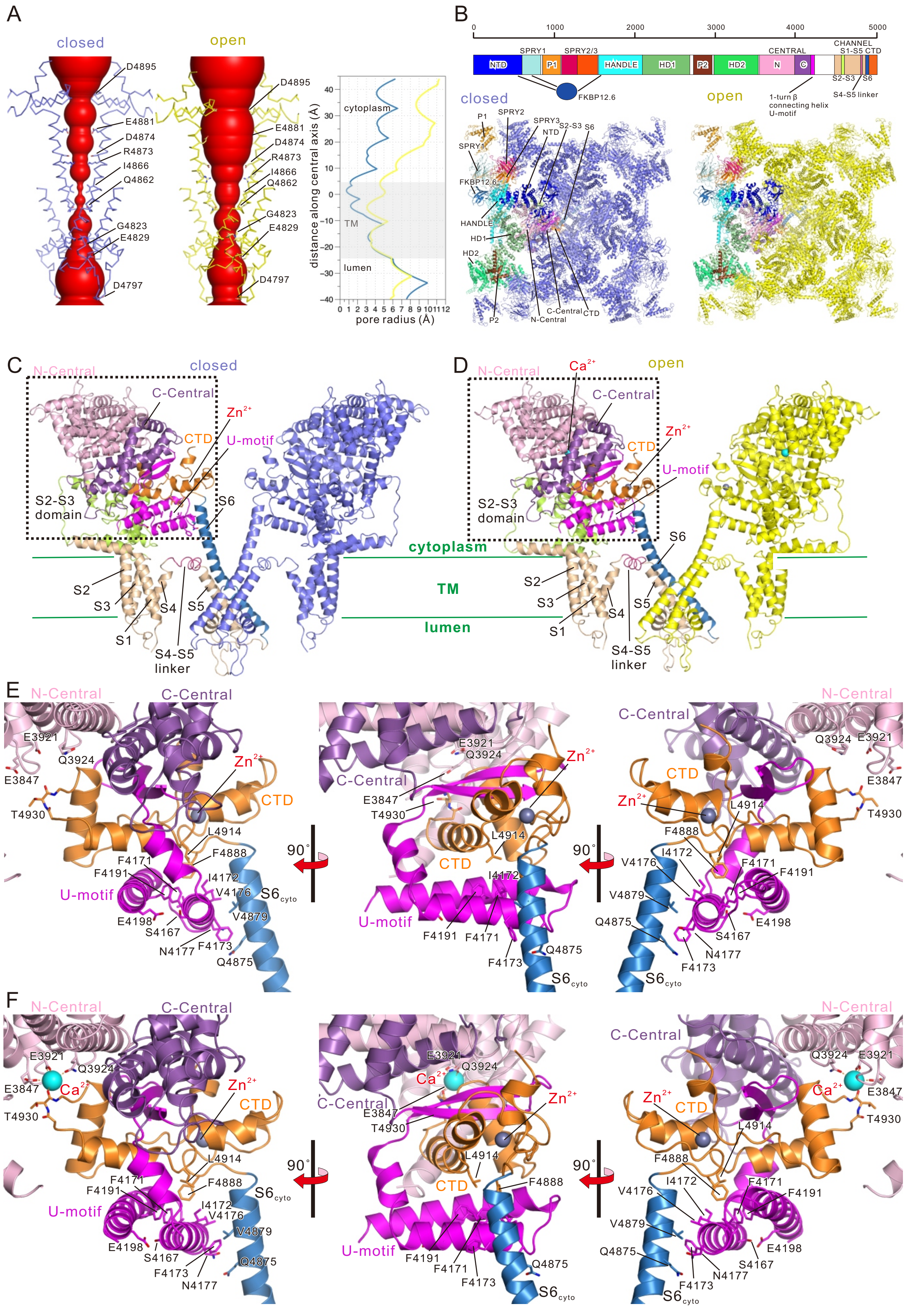
A
closed
open
D4895
E4881
D4874
R4873
I4866
Q4862
G4823
E4829
D4797
cytoplasm
TM
lumen
distance along central axis (Å)
pore radius (Å)
B
SPRY1
SPRY2/3
HANDLE
HD1
HD2
CENTRAL
CHANNEL
S1-S5 CTD
NTD
P1
P2
N
C
S2-S3
S6
S4-S5 linker
FKBP12.6
1-turn β
connecting helix
U-motif
C
N-Central
C-Central
Zn2+
CTD
U-motif
S2-S3
domain
S6
cytoplasm
TM
lumen
S1
S2
S3
S4
S5
S4-S5
linker
D
Ca2+
E
F
E3921
Q3924
E3847
T4930
L4914
F4888
F4171
F4191
I4172
V4176
V4879
Q4875
E4198
S4167
N4177
F4173
90°
Figure S2

Figure S2

**Figure. S2. Detail of the structures of RyR2, related to Figure 1.**

(A) Channel pore and pore radii along the ion conducting pathway of the closed and open states of RyR2 calculated by HOLE (Smart et al., 1993).

(B) (Top) Domain organization of rat RyR2 protomer. (Bottom) Structures of RyR2 viewed from cytoplasmic side (top view) in the closed state (left) and open state (right). One of four protomers in each state is colored, and the color of each domain follows the color shown in (top).

(C and D) Ribbon model of core domains of RyR2 in the closed (C) and open (D) state. In the left protomer, each domain is colored (N-Central, light pink; C-Central, purple; U-motif, magenta; S1-S5, wheat; S2-S3 domain, light green; S4-S5 linker, warm pink; S6, blue; CTD, orange). $Ca^{2+}$, shown as cyan ball; $Zn^{2+}$, shown as gray ball.

(E and F) Magnified views of the dotted box in (C) or (D), respectively. Only N-/C-Central, U-motif, $S6_{cyto}$, and CTD are shown. The middle and right were rotated 90º and 180º to the left, respectively. $Ca^{2+}$-binding site is composed of the carbonyl oxygen of T4930 (CTD) from the lower side and the side chains (E3847, E3921, and Q3924) from N-Central.

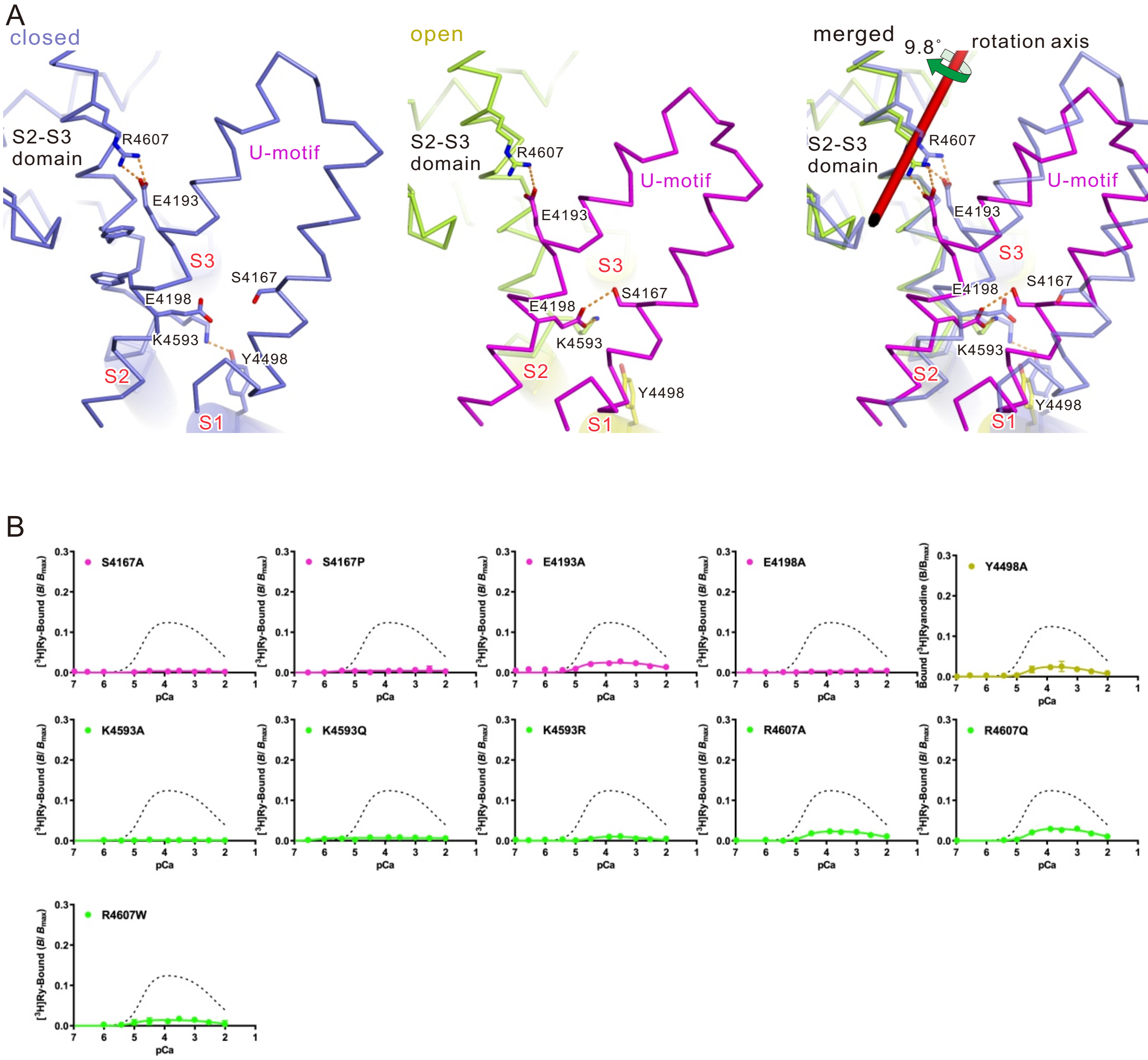
A
closed
open
merged
9.8°
rotation axis
S2-S3 domain
R4607
E4193
U-motif
S3
S4167
E4198
K4593
Y4498
S2
S1
B
S4167A
S4167P
E4193A
E4198A
Y4498A
K4593A
K4593Q
K4593R
R4607A
R4607Q
R4607W
[3H]Ry-Bound (B/ Bmax)
Bound [3H]Ryanodine (B/Bmax)
pCa

Figure S3

**Figure. S3. Movement of the S2-S3 domain and functional analysis of the related mutants, related to Figure 2.**

(A) Interface of the U-motif and S2-S3 domain in the closed state (left), open state (middle), and overlay of both states (right) shown as a Cα model viewed from cytoplasmic side. Amino acid residues involved in the key interactions are shown as stick models. The color of carbon atoms is the same as that of Cα; oxygen, red; nitrogen, blue. The TM region is indicated in yellow and the region forming α-helices is overlaid with the cylinder model. Hydrogen bonds are shown as orange dotted lines.

(B) Functional analysis of mutants involved in the U-motif/S2S3 domain interaction. $Ca^{2+}$-dependent [$^3$H]ryanodine binding of WT (dotted line) and individual mutants.

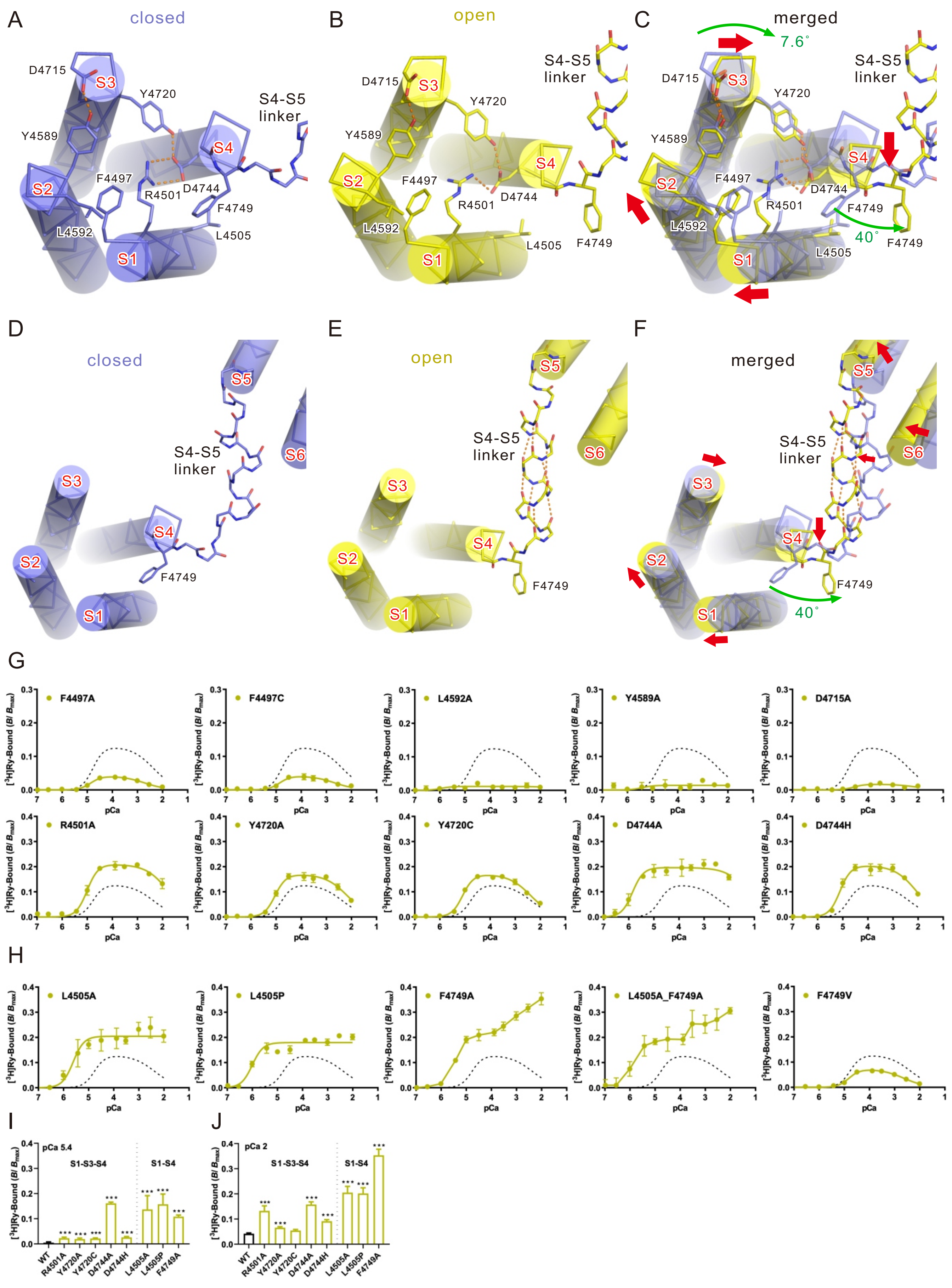
A
closed
B
open
C
merged
7.6°
D4715
S3
Y4720
S4-S5 linker
Y4589
S4
S2
F4497
R4501
D4744
F4749
L4592
L4505
S1
40°
D
closed
E
open
F
merged
S5
S6
S3
S4
S2
S1
F4749
G
F4497A
F4497C
L4592A
Y4589A
D4715A
R4501A
Y4720A
Y4720C
D4744A
D4744H
[3H]Ry-Bound (B/ Bmax)
pCa
H
L4505A
L4505P
F4749A
L4505A_F4749A
F4749V
I
pCa 5.4
S1-S3-S4
S1-S4
WT
R4501A
Y4720A
Y4720C
D4744A
D4744H
L4505A
L4505P
F4749A
J
pCa 2

Figure S4

**Figure. S4. Key interactions in the transmembrane region and functional analysis of the related mutants, related to Figure 3.**

(A–C) The S1-S4 bundle and part of S4-S5 linker. Closed state (A), open state (B), and overlay of the structures in both states (C) are shown as Cα models and overlaid with cylinder models. Viewed from the cytoplasm perpendicular to the membrane. Hydrogen bonds/salt bridges are shown as orange botted lines.

(D-F) Drastic change in the S4-S5 linker upon binding of $Ca^{2+}$. The TM region in the closed (D) and open (E) states, and overlay of the structures in both states (F), viewed from the cytoplasm perpendicular to the membrane, are shown as Cα models and overlaid with cylinder models. The S4-S5 linker did not form an α-helix in the closed state (D), but formed an α-helix in the closed state (E). The color of carbon atoms is the same as that of Cα; oxygen, red; nitrogen, blue. Hydrogen bonds are shown as orange dotted lines.

(G and H) Functional analysis of mutants involved in the movement of the TM region. $Ca^{2+}$-dependent [$^{3}$H]ryanodine binding of WT (dotted line) and individual mutants involved in the S1/S2, S2/S3, S1/S3/S4 (G), or S1-S4 (H) interactions.

(I and J) [$^{3}$H]Ryanodine binding of WT and all mutants at pCa 5.4 (I) and pCa 2 (J). ***$p < 0.001$.

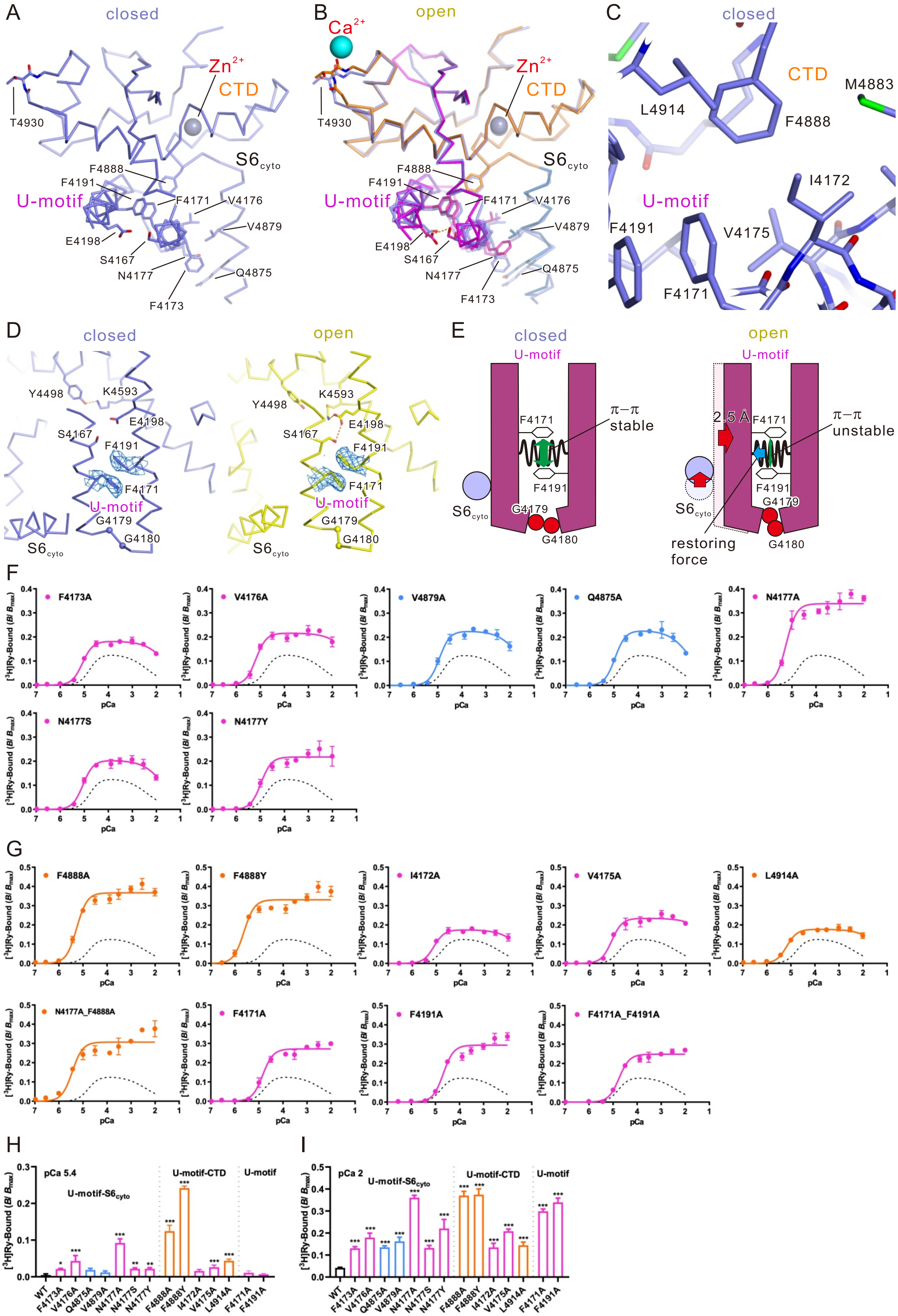

A
closed
Zn2+
CTD
T4930
S6cyto
F4888
F4191
U-motif
F4171
V4176
V4879
E4198
S4167
N4177
Q4875
F4173
B
Ca2+
open
C
M4883
L4914
F4888
I4172
F4191
V4175
F4171
D
Y4498
K4593
E4198
S4167
G4179
G4180
E
π−π
stable
unstable
2.5 Å
restoring force
F
G
H
I
pCa 5.4
pCa 2
U-motif-S6cyto
U-motif-CTD
[3H]Ry-Bound (B/ Bmax)
pCa
F4173A
V4176A
V4879A
Q4875A
N4177A
N4177S
N4177Y
F4888A
F4888Y
I4172A
V4175A
L4914A
N4177A_F4888A
F4171A
F4191A
F4171A_F4191A
WT


Figure S5

**Figure. S5. Key interactions and functional analysis of the related mutants between the U-motif and $S6_{cyto}$/CTD, related to Figures 4.**

(A and B) Structure around the U-motif in the closed state (A) and overlayed with the structure in the open state (B) (U-motif, magenta; $S6_{cyto}$, blue; CTD, orange). $Ca^{2+}$ and $Zn^{2+}$ are shown as cyan and gray spheres, respectively.

(C) Details of the U-motif/CTD interaction around F4888 in the closed state.

(D) Density maps around the side chains of F4171 and F4191 in the closed state and open state contoured at 0.03.

(E) Scheme of the compaction in U-motif and the critical role in the Interaction between F4171/F4191.

(F and G) $Ca^{2+}$-dependent [$^3$H]ryanodine binding of WT (dotted line) and individual mutants involved in the U-motif/$S6_{cyto}$ (F) and U-motif/CTD (G) interactions. (H and I) [$^3$H]Ryanodine binding of WT and all mutants at pCa 5.4 (H) and pCa 2 (I). Data are presented as the mean ± SD. *p < 0.05; **p < 0.01; ***p < 0.001.

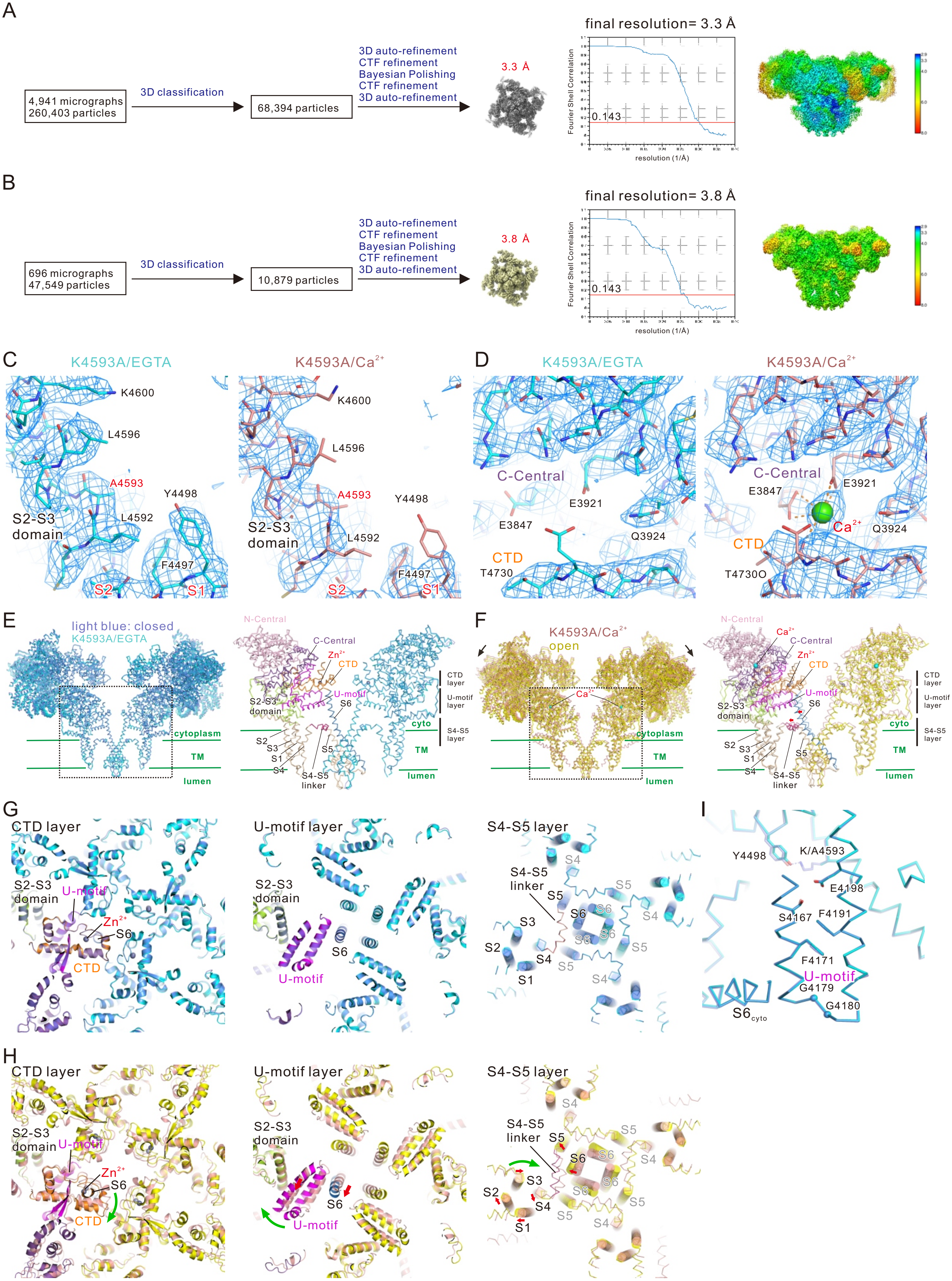

A
4,941 micrographs
260,403 particles
3D classification
68,394 particles
3D auto-refinement
CTF refinement
Bayesian Polishing
CTF refinement
3D auto-refinement
3.3 Å
final resolution= 3.3 Å
Fourier Shell Correlation
0.143
resolution (1/Å)
B
696 micrographs
47,549 particles
3D classification
10,879 particles
3D auto-refinement
CTF refinement
Bayesian Polishing
CTF refinement
3D auto-refinement
3.8 Å
final resolution= 3.8 Å
Fourier Shell Correlation
0.143
resolution (1/Å)
C
K4593A/EGTA
K4593A/Ca2+
K4600
L4596
A4593
Y4498
L4592
S2-S3 domain
F4497
S2
S1
D
C-Central
E3921
E3847
Q3924
CTD
T4730
Ca2+
T4730O
E
light blue: closed
K4593A/EGTA
N-Central
C-Central
Zn2+
CTD
U-motif
S6
S2-S3 domain
CTD layer
U-motif layer
S4-S5 layer
cytoplasm
cyto
TM
lumen
S2
S3
S1
S4
S5
S4-S5 linker
F
K4593A/Ca2+
open
Ca2+
G
CTD layer
U-motif layer
S4-S5 layer
S2-S3 domain
U-motif
Zn2+
S6
CTD
S4-S5 linker
S1
S2
S3
S4
S5
H
I
Y4498
K/A4593
E4198
S4167
F4191
F4171
U-motif
G4179
G4180
S6cyto


Figure S6

**Figure. S6. Single particle analysis on the K4593A mutant, related to Figure 5.**

(A and B) Workflows for cryo-EM data processing and estimation of resolution and local resolution EM map of K4593A mutant in the presence of EGTA (K4593A(EGTA)) (A) and in the presence of $Ca^{2+}$ (K4593A($Ca^{2+}$)) (B).

(C and D) Density maps for the reconstructed structures of K4593A(EGTA)and K4593A($Ca^{2+}$), respectively. Around mutated residue (A4593) (C) and the $Ca^{2+}$ binding site (D), contour level at 0.035.

(E) Overlay of WT in the closed state shown in light blue and K4593A(EGTA) shown in cyan (left) and the magnified view of the dotted box in the left (right).

(F) Overlay of K4593A($Ca^{2+}$) shown in salmon and WT in the open state shown in yellow (left) and the magnified view of the dotted box in the left (right).

(G and H) Cross-section views of CTD, U-motif, and S4-S5 layers. WT in the closed state is colored with light blue and K4593A(EGTA) is colored according to Figure 1B or cyan (G). K4593A($Ca^{2+}$) is colored with salmon and WT in the open state is colored according to Figure 1B or yellow (H).

(I) Analysis of the compactization in U-motif. Overlay of the structure in the closed state (light blue) and K4593A(EGTA) (cyan) are shown as a Cα model. The structures are fitted in the C-terminal side of U-motif (4183-4205).

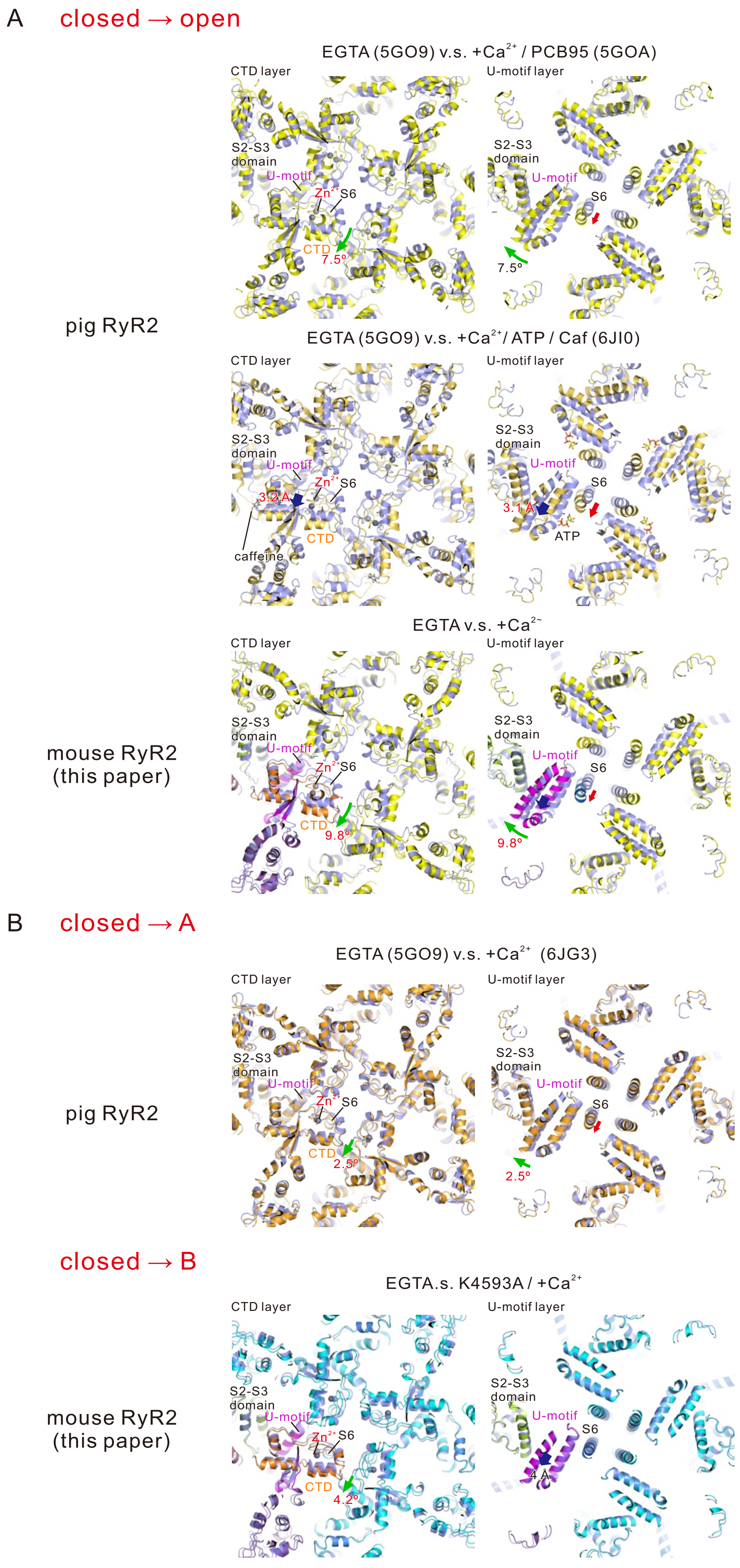
A
closed → open
EGTA (5GO9) v.s. +Ca2+ / PCB95 (5GOA)
CTD layer
U-motif layer
S2-S3 domain
U-motif
Zn2+
S6
CTD
7.5°
pig RyR2
EGTA (5GO9) v.s. +Ca2+/ ATP / Caf (6JI0)
3.2 A
3.1 A
caffeine
ATP
EGTA v.s. +Ca2+
mouse RyR2 (this paper)
9.8°
B
closed → A
EGTA (5GO9) v.s. +Ca2+ (6JG3)
2.5°
closed → B
EGTA.s. K4593A / +Ca2+
4.2°
4 A

Figure S7

**Figure. S7. Structural changes detected in previously reported structures.**

(A) Cross-section views of CTD and U-motif layers of pig RyR2 and mouse RyR2. (1st and 2nd rows) Two different structures in the open state (5GOA and 6JIO) overlayed with the structure in the closed state (5GO9). The colors are as follows; in the presence of EGTA (5GO9), light blue; in the presence of $Ca^{2+}$ and PCB95 (5GOA), yellow; in the presence of $Ca^{2+}$, ATP, and caffein (6JI0). (3rd row) The structure of mouse RyR2 in the open state overlayed with the structure in the closed state. Closed state is colored in light blue and open state is colored according to Figure 1B or yellow.

(B) Cross-section views of CTD and U-motif layers of pig RyR2 and mouse RyR2. (4th row) The structure of pig RyR2 in the presence of $Ca^{2+}$ overlayed with the structure in the closed state. The colors of pig RyR2 are as follows; in the presence of EGTA (5GO9), light blue; in the presence of $Ca^{2+}$ (6JG3), orange. (5th row) The structure of mouse K4593A($Ca^{2+}$) overlayed with the structure of mouse K4593A(EGTA). K4593A(EGTA) is colored with cyan and K4593A($Ca^{2+}$) is colored according to Figure 1B or salmon.

**Table S1. Data collection, rotamer outliers, processing, model refinement, and validation**

| Protein | WT | | | | | | K4593A | |
|---|---|---|---|---|---|---|---|---|
| Condition | EGTA | | | 100 µM $Ca^{2+}$ | | | EGTA | 100 µM $Ca^{2+}$ |
| State | before classification | class1 | class2 | class1 | class2 | class3 | | |
| PBD ID | 7VML | 7VMM | 7VMN | 7VMO | 7VMP | 7VMQ | 7VMR | 7VMS |
| EMDB ID | EMD-30688 | EMD-30689 | EMD-30690 | EMD-30691 | EMD-30692 | EMD-30693 | EMD-32036 | EMD-32037 |
| **Data collection and processi** | | | | | | | | |
| Microscope | Titan Krios G3i | | | | | | | |
| Detector | Gatan K3 BioQuantum | | | | | | | |
| Magnification | x81k | | | x105k | | | x105k | x105k |
| Voltage (kV) | 300 | | | | | | | |
| Electron exposure ($e^-$/Å$^2$) | 50 | | | 60 | | | 50 | 50 |
| Defocus range (µm) | 0.5 - 2.0 | | | | | | | |
| Pixcel size (Å) | 1.07 | | | 0.83 | | | 0.83 | |
| Symmetry imposed | *C* 4 | | | | | | | |
| Initial particle images | 578,868 | | | 388,457 | | | 260,403 | 47,549 |
| Final particle images | 109,425 | 45,120 | 41,197 | 45,432 | 42,375 | 40,665 | 68,394 | 10,879 |
| Map resolution (Å) | 3.3 | 3.49 | 3.52 | 3.45 | 3.51 | 3.72 | 3.3 | 3.8 |
| FSC threshold | 0.143 | 0.143 | 0.143 | 0.143 | 0.143 | 0.143 | 0.143 | 0.143 |
| Map sharpening B-factor | -67 | -22 | -22 | -49 | -46 | -65 | -50 | -84 |
| **Model building and refineme** | | | | | | | | |
| Model composition | | | | | | | | |
| Protein atoms | 123564 | 123564 | 123564 | 122032 | 122032 | 122032 | 123548 | 123548 |
| Metals | 4 | 4 | 4 | 8 | 8 | 8 | 4 | 8 |
| R.M.S. deviations | | | | | | | | |
| Bond length (Å) | 0.002 | 0.002 | 0.003 | 0.002 | 0.003 | 0.003 | 0.003 | 0.003 |
| Bond angles (°) | 0.502 | 0.527 | 0.566 | 0.481 | 0.596 | 0.645 | 0.661 | 0.644 |
| **Validation** | | | | | | | | |
| MolProbity score | 1.96 | 2.36 | 2.25 | 2.24 | 2.5 | 2.63 | 2.18 | 2.03 |
| Clashscore | 8.17 | 8.32 | 9.02 | 8.59 | 10.32 | 11.08 | 10.77 | 12.85 |
| Rotamer outliers (%) | 1.39 | 4.51 | 2.87 | 2.88 | 4.95 | 6.61 | 1.87 | 0.14 |
| Ramachandran plot | | | | | | | | |
| Favored (%) | 93.85 | 93.8 | 93.71 | 93.37 | 93.18 | 93.14 | 93.59 | 93.8 |
| Allowed (%) | 6.15 | 6.2 | 6.29 | 6.63 | 6.82 | 6.86 | 6.41 | 6.2 |
| Outlier (%) | 0 | 0 | 0 | 0 | 0 | 0 | 0 | 0 |

Clashscores, rotamer outlers, and Ramachandran plots were calculated using PHENIX (Adams et al., 2010; Afonine et al., 2018).
